# The sole essential low molecular weight tropomyosin isoform of *Caenorhabditis elegans* is essential for pharyngeal muscle function

**DOI:** 10.1101/2024.12.13.628433

**Authors:** Michael J. Kimmich, Meaghan A. Geary, Lei Mi-Mi, SarahBeth D. Votra, Christopher D. Pellenz, Sumana Sundaramurthy, David Pruyne

## Abstract

Tropomyosin is an actin-binding protein that plays roles ranging from regulating muscle contraction to controlling cytokinesis and cell migration. The simple nematode *Caenorhabditis elegans* provides a useful model for studying the core functions of tropomyosin in an animal, having a relatively simple anatomy, and a single tropomyosin gene, *lev-11*, that produces seven isoforms. Three higher molecular weight isoforms (LEV-11A, D, O) regulate contraction of body wall and other muscles, but comparatively less is known of the functions of four lower molecular weight isoforms (LEV-11C, E, T, U). We demonstrate here *C. elegans* can survive with a single low molecular weight isoform, LEV-11E. Mutants disrupted for LEV-11E die as young larvae, whereas mutants disrupted for all other short isoforms are viable with no overt phenotype. Vertebrate low molecular weight tropomyosins are often considered “nonmuscle” isoforms, but we find LEV-11E localizes to sarcomeric thin filaments in pharyngeal muscle, and co-precipitates from worm extracts with the formin FHOD-1, which is also associated with thin filaments in pharyngeal muscle. Pharyngeal sarcomere organization is grossly normal in larvae lacking LEV-11E, indicating the tropomyosin is not required to stabilize thin filaments, but pharyngeal pumping is absent, suggesting LEV-11E regulates actomyosin activity similar to higher molecular weight sarcomeric tropomyosin isoforms.

## INTRODUCTION

In the sarcomeres of striated muscle, actin-based thin filaments interdigitate with and slide past myosin-based thick filaments to give rise to contractile force (Huxley & Hanson, 1954). An important aspect of this mechanism is the precise regulation of contraction. Tropomyosin (Tpm) and the troponin complex are crucial to regulating the binding of myosin to thin filaments in striated muscle (Gunning et al., 2015). Tpm forms coiled-coil dimers that bind adjacent Tpm dimers in a head-to-tail fashion as they decorate actin filaments. In relaxed muscle, Tpm blocks binding of muscle myosin to thin filaments, preventing contraction. With muscle activation, Ca^2+^ induces a conformational change in the troponin complex, which shifts the position of Tpm to expose myosin-binding sites on thin filaments, permitting contraction. Outside its role in regulating muscle contraction, the presence of Tpm on actin filaments can slow depolymerization at pointed ends, block filament severing by actin-depolymerization factor (ADF)/cofilin or gelsolin, alter actin filament stiffness, differentially regulate the binding of myosin isoforms, and stimulate formin-dependent actin polymerization (Phillips et al., 1986; Broschat et al., 1989; Goldmann, 2000; Gordon et al., 2000; Ono & Ono, 2002; Wawro et al., 2007; Alioto et al., 2016; Bicskei et al., 2018). These activities likely contribute to the roles Tpm plays in non-muscle cells in regulating cell migration, modulating stress fiber formation, and supporting cytokinesis (Thoms et al., 2008; Tojkander et al., 2011; Lees et al., 2013).

Mammals have four Tpm genes that generate up to 40 isoforms through alternative splicing or utilizing different promoters (Gunning et al., 2015). Broadly speaking, these isoforms are sub-divided into higher molecular weight (long) and lower molecular weight (short) isoforms. Messages for long isoforms initiate from a promoter upstream of the first exon, and utilize exons 1a and either 2a or 2b of their respective Tpm gene, while short isoforms utilize a promoter downstream of exons 2a/2b to initiate transcription at exon 1b (Geeves et al., 2015). All mammalian Tpm isoforms that associate with thin filaments in muscle sarcomeres (so-called sarcomeric Tpm isoforms) are long isoforms, whereas Tpms expressed in nonmuscle cells or associated with non-sarcomeric actin filaments in muscle (so-called cytosolic Tpm isoforms) include long and short isoforms (Hardeman et al., 2020).

The simple nematode *C. elegans* represents a conserved but relatively simpler model for studying Tpm function. The worm genome encodes one Tpm gene, *lev-11*, which gives rise to seven isoforms through use of two promoters and alternative splicing (Anyanful et al., 2001; Barnes et al., 2018; Watabe et al., 2018; Ono et al., 2023) (Fig.1). Similar to mammalian Tpm genes, initiation of transcription from an early promoter in *lev-11* generates long Tpm isoforms (LEV-11A, D, and O) that utilize exons 1, 2, and 3a, whereas transcription from a promoter downstream of exon 3a generates short isoforms (LEV-11C, E, T, and U) that utilize exon 3b as their starting exon (Ono et al., 2023). Also similar to mammals, the long isoforms LEV-11A, D, and O associate with sarcomeric thin filaments in the striated body wall muscle, where they regulate contraction and thin filament stability (Kagawa et al., 1995; Barnes et al., 2018; Ono & Ono, 2002). Little is known of the functions of the short Tpm isoforms of *C. elegans*. Analysis of minigene reporters for alternative splicing has shown exon selection is tissue-specific, resulting in LEV-11C being expressed primarily in the intestine and neurons, LEV-11E being primarily expressed in the pharynx and excretory cell, while the expression patterns for LEV-11T and U remain to be determined (Anyanful et al., 2001; Watabe et al., 2018; Ono et al., 2023). Induction of RNA interference (RNAi) targeting *lev-11* exons 3b and 4a (common to all short isoforms) in the adult germline resulted in 70-75% of progeny experiencing embryonic arrest at the 300-cell stage, while 15% of surviving larvae had deformed pharynges, suggesting short Tpm isoforms are essential for early development in *C. elegans* (Anyanful et al., 2001). RNAi targeting exon 5c, which is utilized exclusively by LEV-11C, induced similar phenotypes, suggesting LEV-11C is a key essential isoform (Anyanful et al., 2001). In contrast, a CRISPR-based knockout targeting exon 7c unique for LEV-11U resulted in only modest motility defects and reduced brood size (Ono et al., 2023), suggesting relatively minor roles for that isoform.

**Figure 1.**
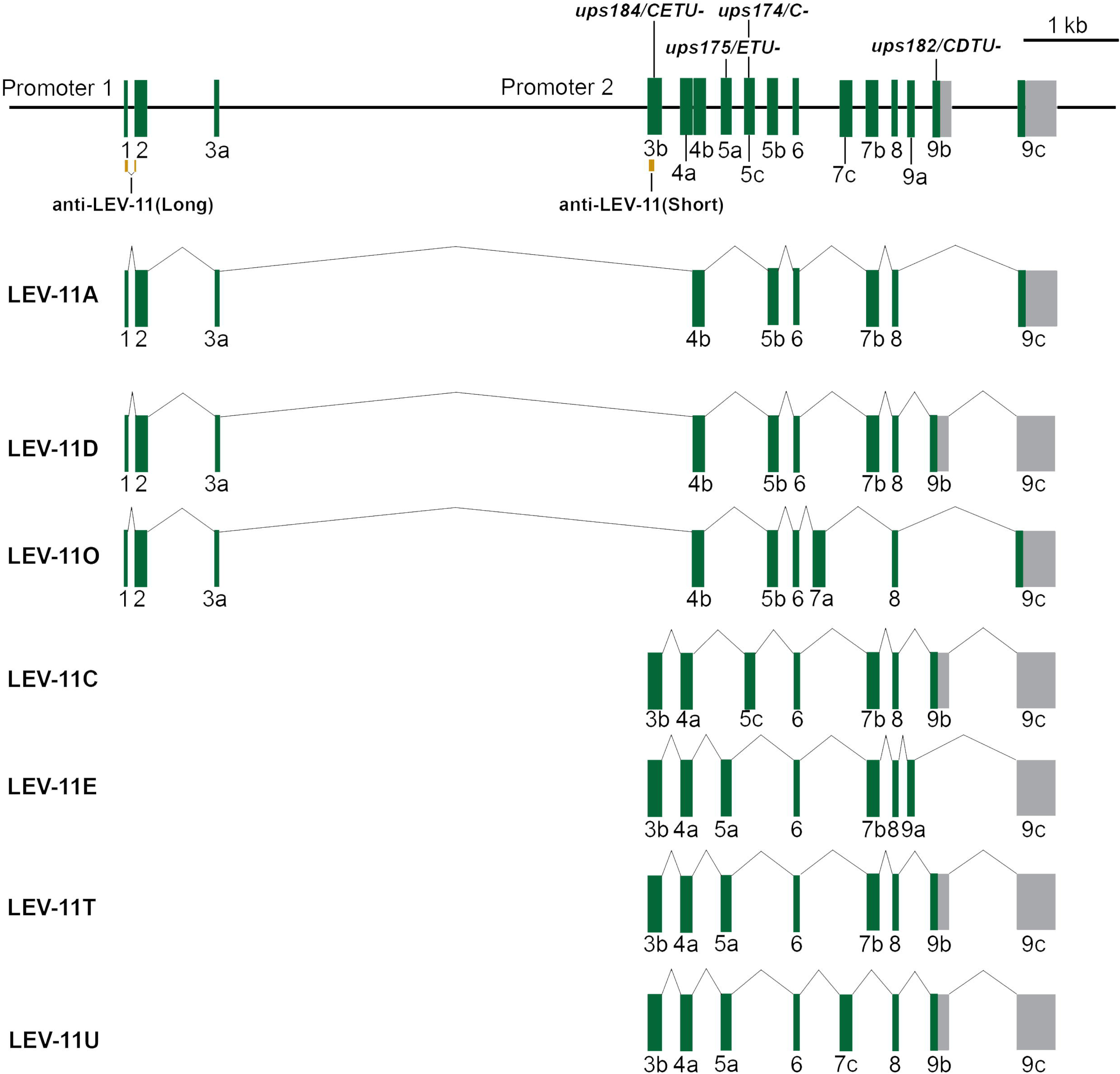
Structure of the *C. elegans* tropomyosin gene, *lev-11*, and exon usage for LEV-11 splice isoforms. Exons are numbered and shown with protein-coding sequences in green and non-coding sequences in grey. At top are indicated positions of nonsense mutation alleles generated during this study, and the location of sequences encoding peptides used to generate anti-LEV-11(Long) and anti-LEV-11(Short) in orange. Transcripts for long LEV-11 isoforms (A, D, and O) initiate from Promoter 1 upstream of Exon 1, while transcripts for short LEV-11 isoforms (C, E, T, and U) initiate from Promoter 2 located between exons 3a and 3b. Scale bar, 1 kb.

Using CRISPR-based mutations to disrupt single exons in the *lev-11* gene, we demonstrate here that LEV-11C and T, like LEV-11U, are not required for viability in *C. elegans*, but animals lacking LEV-11E expression in pharyngeal muscle exhibit pharyngeal muscle paralysis and die during early larval development. LEV-11E associates with sarcomeric thin filaments in pharyngeal muscle, suggesting this short isoform functions as a sarcomeric Tpm.

## RESULTS

### FHOD-1 co-precipitates with one or more short LEV-11 isoforms

The formin FHOD-1 is a putative actin nucleating factor expressed in all muscle cells in *C. elegans*, including pharyngeal muscle used for feeding, and body wall muscle used for crawling and swimming (Mi-Mi et al., 2012; Mi-Mi & Pruyne, 2015; Sundaramurthy et al., 2020; Kimmich et al., 2024). In order to identify potential binding partners for FHOD-1 in body wall muscle, we performed immunoprecipitation (IP) followed by mass spectrometry (MS) on FHOD-1 from worms that either express a functional GFP-tagged FHOD-1 (FHOD-1::GFP) at an ectopic locus (and thus overexpress FHOD-1), or that bear the putative null deletion allele *fhod-1(tm2363)*, referred to here as *fhod-1(*Δ). Whole worm lysates were isolated from asynchronous populations as well as from worms synchronized at late larval stages L3 to L4, and fractionated to enrich for body wall muscle sarcomere proteins before subjecting to IP using an antibody raised against the FHOD-1 FH2 domain (Mi-Mi et al., 2012). Based on total spectrum counts after MS analysis of pre-IP extracts, actin and body wall muscle myosin II heavy chain UNC-54/MYO-4 were the most abundant proteins, suggesting an enrichment for body wall muscle sarcomere proteins (Supplementary Table S1). However, an abundance of intestine-expressed vitellogenins and pharyngeal muscle-specific myosin II heavy chains MYO-1 and MYO-2 demonstrated significant contamination from other tissues. FHOD-1 was not detected in pre-IP extracts, suggesting the formin is of low abundance, but was strongly enriched after FHOD-1 IP of extracts of FHOD-1-over-expressing but not *fhod-1(*Δ) worms (Supplementary Table S2, Supplementary Table S3). Sarcomere proteins were enriched in FHOD-1 IP samples from FHOD-1-overexpressing and from *fhod-1(*Δ) extracts (Table 1), indicating there was significant non-specific precipitation of those proteins. Despite this high background, we observed short isoforms of Tpm/LEV-11 were recovered after FHOD-1 IP only from extracts of FHOD-1-overexpressing worms. This was true for extracts from asynchronous worms and from late-stage larvae (Table 1). MS analysis showed similar amounts of short Tpm1/LEV-11 isoforms in pre-IP extracts of *fhod-1(*Δ) and FHOD-1-over-expressing animals (Table 1), suggesting their specific recovery from FHOD-1-overexpressing worm extracts reflects co-precipitation with FHOD-1, rather than reduced expression in *fhod-1(*Δ) animals.

**Table 1.** Cytoskeleton-associated proteins detected in extracts and FHOD-1 IP.

| protein (alternative name)<sup>†</sup> | IP from late larvae extract<sup>‡</sup> |  | IP from mixed age worm extract<sup>‡</sup> |  | mixed age worm extract prior to IP<sup>‡</sup> |  |
| --- | --- | --- | --- | --- | --- | --- |
|  | *fhod-1(+)*<sup>§</sup> | *fhod-1(Δ)*<sup>§</sup> | *fhod-1(+)*<sup>§</sup> | *fhod-1(Δ)*<sup>§</sup> | *fhod-1(+)*<sup>§</sup> | *fhod-1(Δ)*<sup>§</sup> |
| UNC-54 (Muscle Myosin) | 418 | 169 | 347 | 439 | 403 | 354 |
| ACT-1/3 (Actin) | 190 | 45 | ND | ND | 419 | 269 |
| MYO-2 (Muscle Myosin) | 150 | 24 | 197 | 244 | 217 | 206 |
| **FHOD-1 (Formactin)** | **148** | **0** | **135** | **0** | **ND** | **ND** |
| LET-75 (Muscle Myosin) | 132 | 16 | 159 | 195 | 176 | 157 |
| MYO-3 (Muscle Myosin) | 124 | 25 | 128 | 179 | 147 | 128 |
| LEV-11A/B/D/F (Tropomyosin)<sup>¶</sup> | 95 | 14 | 32 | 30 | 29 | 28 |
| ACT-5 (Actin) | 65 | 24 | 50 | 45 | 63 | 50 |
| T25F10 (Click-1) | 58 | 8 | 30 | 29 | 29 | 22 |
| MYO-5 (Muscle Myosin) | 55 | 8 | 77 | 126 | 67 | 54 |
| **LEV-11C/E (Tropomyosin)** | **54** | **0** | **21** | **0** | **11** | **14** |
| UNC-15 (Paramyosin) | 52 | 37 | 40 | 42 | 122 | 142 |
| UNC-87 (Calponin-Related) | 35 | 7 | 27 | 25 | 23 | 16 |
| TNT-2 (Troponin T) | 28 | 5 | 15 | 19 | 15 | 8 |
| MUP-2 (Troponin T) | 26 | 6 | 20 | 17 | 14 | 11 |
| MLC-1 (Myosin Regulatory Light Chain) | 21 | 4 | ND | ND | 32 | 30 |
| MLC-3 (Myosin Essential Light Chain) | 16 | 0 | 22 | 13 | 19 | 22 |
| TBB-2 (Beta Tubulin) | 14 | 21 | 31 | 33 | 26 | 24 |
| UNC-27 (Troponin I) | 11 | 0 | 12 | 6 | 12 | 5 |
| TBA-2 (Alpha Tubulin) | 7 | 11 | 10 | 15 | 21 | 19 |
| NMY-2 (Nonmuscle Myosin) | 6 | 0 | 8 | 31 | 23 | 14 |
| TNI-3 (Troponin I) | 5 | 0 | 0 | 0 | 5 | 0 |
| TNI-1 (Troponin I) | 5 | 0 | 0 | 0 | 4 | 0 |
| IFB-2 (Intermediate Filament) | 3 | 0 | 0 | 8 | 13 | 20 |
| LMN-1 (Lamin) | 0 | 3 | 0 | 5 | 19 | 16 |
| LFI-1 (CROCC) | 0 | 86 | 0 | 0 | 0 | 0 |
| HUM-8 (Class VI Myosin) | 0 | 5 | 0 | 10 | 0 | 0 |
| GSNL-1 (Gelsolin) | 0 | 0 | 0 | 4 | 8 | 11 |
| KETN-1 (Kettin) | 0 | 0 | 0 | 12 | 28 | 31 |
| UNC-94 (Tropomodulin) | 0 | 0 | 3 | 4 | 3 | 0 |
| MYO-6 (Muscle Myosin) | 0 | 0 | 0 | 5 | 6 | 6 |
| ACT-4 (Actin) | 0 | 0 | 126 | 112 | 0 | 0 |
| MLC-2 (Myosin Regulatory Light Chain) | 0 | 0 | 29 | 18 | 27 | 24 |

<sup>†</sup> Proteins were identified using the *Caenorhabditis elegans* UNIPROT database, accessed 11/17/2012. Minimum requirements for protein identification were 99.9% probability of correct protein with 95% correct peptide identification.
<sup>‡</sup> Numbers shown are total spectrum counts.
<sup>§</sup> *fhod-1(+)* represents worms bearing wild type endogenous *fhod-1* plus ectopic *fhod-1::gfp* transgene; *fhod-1(Δ)* indicates worms bearing *fhod-1(tm2363)* putative null allele.
<sup>¶</sup> Isoforms b and f are predicted splice products of *lev-11* but to date there is no evidence demonstrating they are expressed.

### Short LEV-11 isoforms are essential for larval development

To test whether short Tpm/LEV-11 isoforms functionally interact with FHOD-1 *in vivo*, we set out to disrupt their expression. Unfortunately, we were unable to determine from MS analysis which short LEV-11 isoforms precipitate with FHOD-1 due to extensive shared sequences among them. Thus, to monitor expression of all short and all long LEV-11 isoforms, we raised antibodies against two different antigens. Anti-LEV-11(Long) was raised against a peptide coded within *lev-11* exons 1 and 2 (the starting two exons for all long isoforms) and anti-LEV-11(Short) was raised against a peptide coded within exon 3b (the starting exon for all short isoforms) (Fig.1). Based on western blot analysis, these antibodies specifically recognize their respective isoforms. That is, anti-LEV-11(Long) recognized bacterially expressed and purified long isoform LEV-11A but not short isoforms LEV-11C or LEV-11E, while anti-LEV-11(Short) recognized recombinant LEV-11C and E, but not LEV-11A (Fig.2A). Similarly, anti-LEV-11(Long) recognized in whole worm extracts an endogenous protein of approximately the same molecular weight as recombinant LEV-11A, and also recognized exogenously expressed higher molecular weight amino-terminally (N-terminally) GFP-tagged LEV-11A (GFP::LEV-11A), but not GFP::LEV-11C or GFP::LEV-11E (Fig.2A). In contrast, anti-LEV-11(Short) recognized in worm extracts endogenous proteins of similar molecular weight as recombinant LEV-11C and E, and also recognized overexpressed GFP::LEV-11C and GFP::LEV-11E, but not GFP::LEV-11A (Fig.2A).

**Figure 2.**
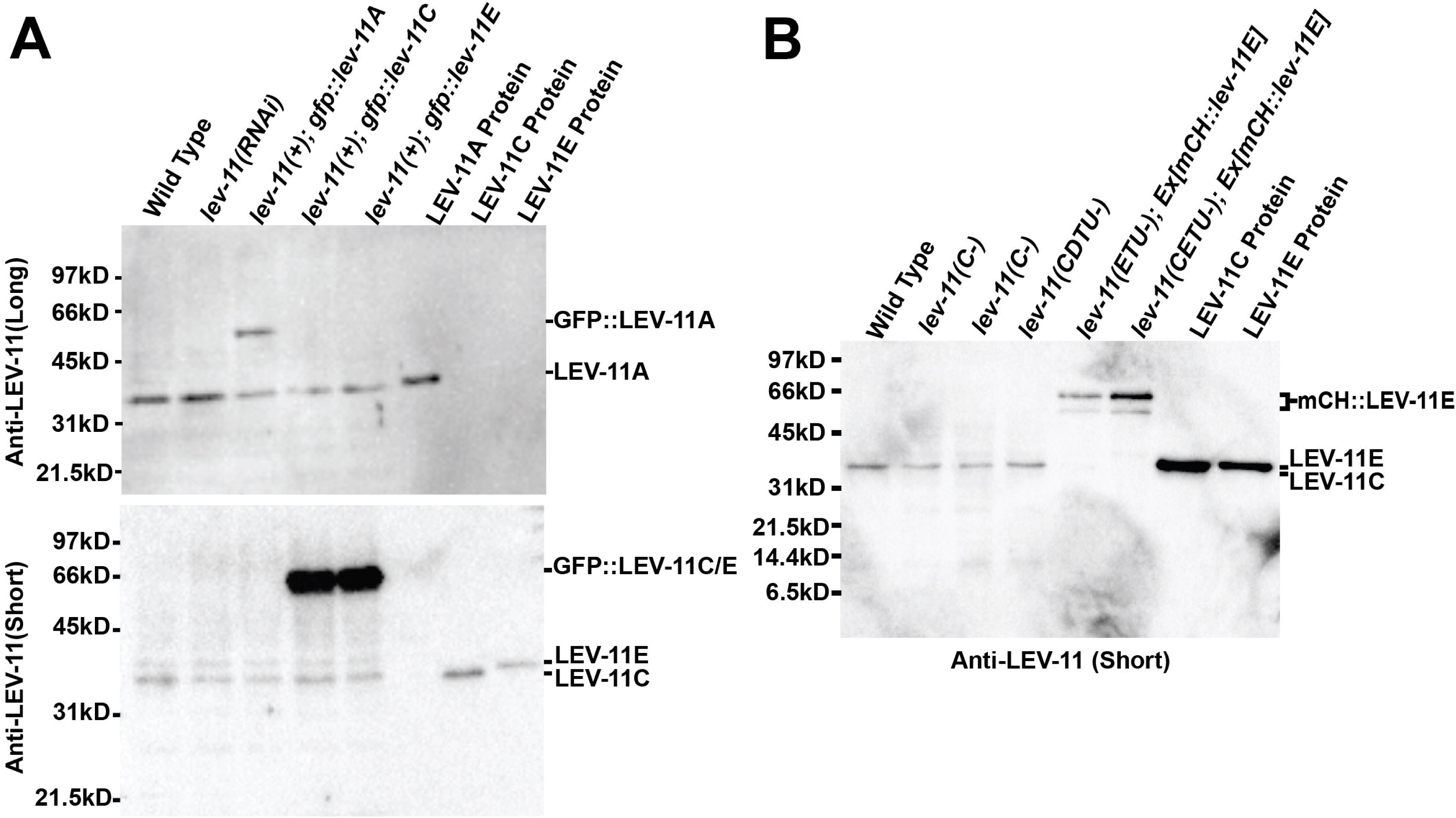
LEV-11 isoform detection and expression. Whole worm extracts (designated by worm genotype) and purified recombinant LEV-11 proteins (long isoform LEV-11A and short isoforms LEV-11C and E) were probed by immunoblot using antibodies specific for long or short isoforms. **(A)** Anti-LEV-11(Long) recognizes purified LEV-11A but not LEV-11C or E, and recognizes endogenous protein of similar molecular weight to LEV-11A, and exogenous GFP-tagged LEV-11A, but not endogenous or GFP-tagged LEV-11C or E. Conversely, anti-LEV-11(Short) recognizes purified LEV-11C and E but not A, and recognizes endogenous proteins of similar molecular weight to LEV-11C/E, and exogenous GFP-tagged LEV-11C and E, but not endogenous or GFP-tagged LEV-11A. **(B)** Immunoblotting whole worm extracts and purified recombinant LEV-11 proteins with anti-LEV-11(Short) shows worms bearing CRISPR-generated mutations simultaneously targeting LEV-11E, T, and U, or simultaneously targeting LEV-11C, E, T, and U lack most or all endogenous low molecular weight LEV-11, but express exogenous mCherry-tagged LEV-11E.

We attempted to knock down expression of all short LEV-11 isoforms using RNAi, simultaneously targeted exon combinations 3a-4a-5a (the first three exons for LEV-11E, T, U) and 3a-4a-5b (the first three exons for LEV-11C). However, western blots probed with anti-LEV-11(Short) and anti-LEV-11(Long) showed no changes in LEV-11 expression (Fig.2A), and we observed no phenotypes among the progeny of treated animals. As an alternative approach, we used CRISPR/Cas9-mediated mutagenesis to insert stop codons into *lev-11* exons utilized by different subsets of LEV-11 isoforms (Fig.1). As previous RNAi-based studies implicated LEV-11C as an essential isoform (Anyanful et al., 2001), we first inserted premature stop codons into exon 5c to generate *lev-11(C-)* alleles disrupted for LEV-11C only (Fig.1). Surprisingly, animals homozygous for any of three independently isolated *lev-11(C-)* alleles were viable and exhibited no reduction in the number of eggs produced (Fig.3A,B). By western blot analysis, *lev-11(C-)* animals still expressed significant amounts of low molecular weight LEV-11(Fig.2B). To test whether LEV-11C might function redundantly with other short isoforms, we used CRISPR/Cas9-mediated mutagenesis to insert premature stop codons into exon 9b, producing *lev-11(CDTU-)* targeting short isoforms LEV-11C, T, and U, and long isoform LEV-11D (Fig.1). As with *lev-11(C-)*, homozygous *lev-11(CDTU-)* animals were viable and appeared grossly normal, and expressed significant amounts of low molecular weight LEV-11 (Fig.2B). This suggests three of the four short Tpm isoforms are dispensable, although we note that the position of the *lev-11(CDTU-)* mutation near the 3’ end of the coding sequences (Fig.1) leaves in question to what degree protein expression or function were affected.

**Figure 3.**
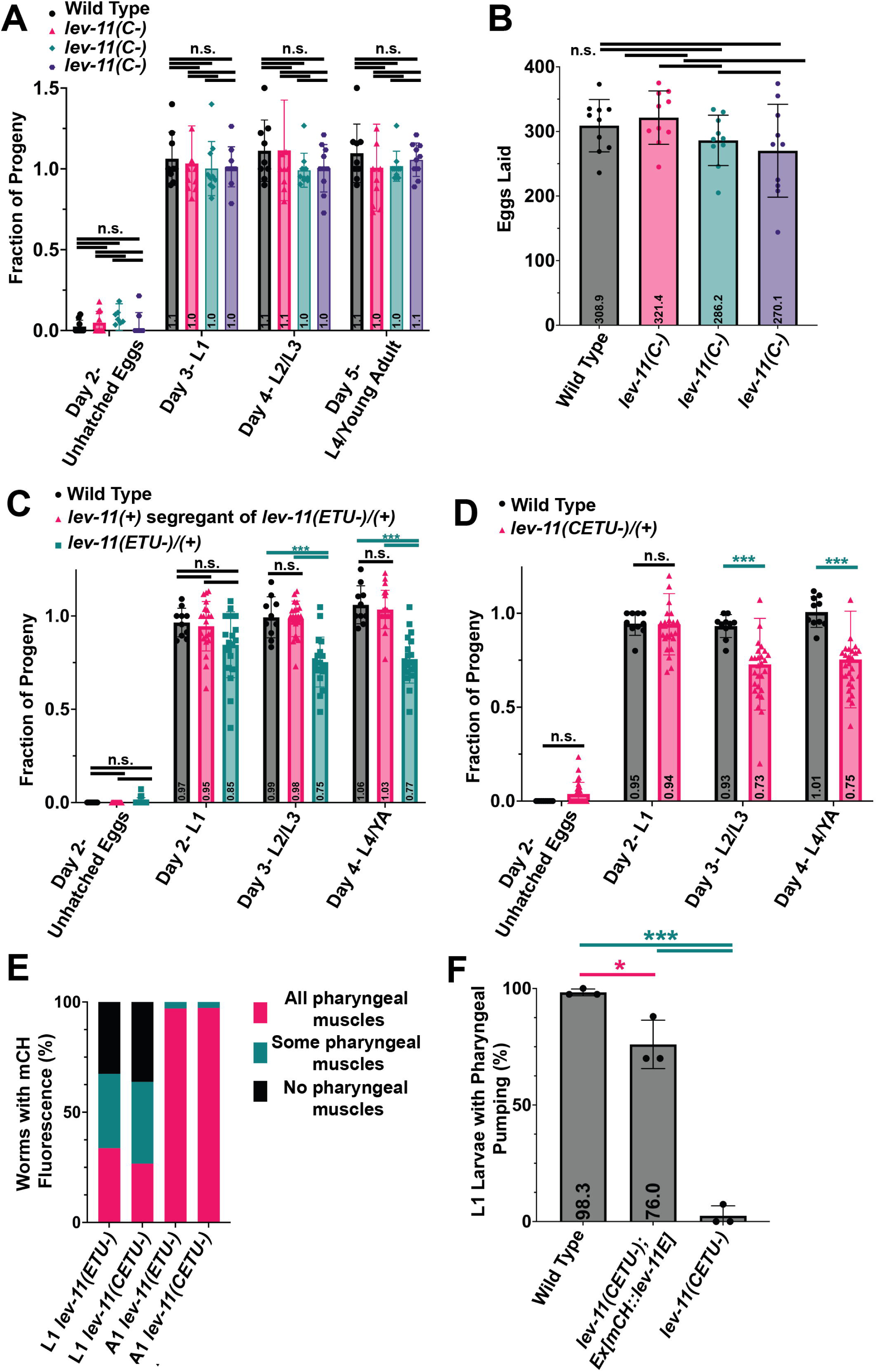
LEV-11E is essential for larval development and pharyngeal muscle function. **(A)** Eggs laid by worms of the indicated genotypes were counted on day 1, and the resultant progeny were examined over 4 days. For worms from three independently isolated *lev-11(C-)* strains, essentially all eggs hatched by day 2, and resultant larvae grew to adulthood (n = 9 wild type animals, 10 animals for each *lev-11(C-)* strain). **(B)** The number of eggs laid was counted over the lifetime of worms of the indicated genotypes, showing *lev-11(C-)* mutants lay the same number of eggs as wild type animals (n = 10 worms per strain). **(C, D)** Viability of progeny was monitored as in **(A)**, showing ∼ 100% of eggs hatch to larvae, but only ∼75% of larvae from **(C)** heterozygous *lev-11(ETU-)/(+)* mutants or **(D)** heterozygous *lev-11(CETU-)/(+)* mutants grew beyond the L1 stage (n = 10 wild type animals, 19 *lev-11(+)* segregants of *lev-11(ETU-)/(+)* animals, 19 *lev-11(ETU-)/(+)* animals in **C**, n = 10 wild type, 18 *lev-11(CETU-)/(+)* animals in **D**). **(E)** *lev-11(ETU-)* and *lev-11(CETU-)* worms with mosaic inheritance of the *Ex[mCH::lev-11E]* ECA were assayed for expression of mCherry-tagged LEV-11E in the pharyngeal muscle of L1 larvae and day 1 adults (A1). In many L1 larvae, some or all pharyngeal muscles lacked mCherry expression, but no day 1 adults lacked mCherry from the entire pharynx, and only a small number (2.9% and 2.7% for *lev-11(ETU-)* and *lev-11(CETU-)*, respectively) lacked mCherry in a small number of pharyngeal muscle cells (n = 86 *lev-11(ETU-)* L1 larvae, 105 *lev-11(CETU-)* L1 larvae, 70 *lev-11(ETU-)* adults, 75 *lev-11(CETU-)* adults). **(F)** Visual observation of wild type or *lev-11(CETU-)* L1 larvae with or without *Ex[mCH::lev-11E]* showed 98% of wild type L1 larvae and 76% of mCH::LEV-11E-expressing larvae exhibit active pharyngeal pumping, while only 2.7% of *lev-11(CETU-)* animals predicted to lack all short LEV-11 isoforms engage in pharyngeal pumping (n = 132 wild type L1 larvae, 89 *lev-11(CETU-); Ex[mCH::lev-11E]* L1 larvae, 85 *lev-11(CETU-)* L1 larvae, over 3 independent trials). (*) *p* = 0.0137; (***) *p* < 0.0001; (n.s.) not significant, *p* ≥ 0.05.

Very different results were obtained when we disrupted expression of the short Tpm isoform, LEV-11E. Exon 9a is exclusive to LEV-11E, but is also positioned near the 3’ end of the coding sequence (Fig.1). Therefore, rather than exclusively targeting LEV-11E, we inserted a missense mutation into exon 5a, predicted to disrupt LEV-11E, T, and U, while sparing LEV-11C (Fig.1). We isolated two independently generated *lev-11(ETU-)* alleles in heterozygous animals, but were unable to recover viable homozygous progeny bearing either allele, suggesting LEV-11E might be essential. We observed that 100% eggs laid by *lev-11(ETU-)/(+)* heterozygotes hatched as L1 larvae, but approximately 25% of these failed to grow beyond the L1 stage (Fig.3C). Genotyping of the surviving adult progeny of *lev-11(ETU-)/(+)* heterozygotes showed all were wild-type or heterozygous for *lev-11(ETU-)*, suggesting *lev-11(ETU-)* homozygotes die during early larval development. In contrast, we observed complete development to adulthood for approximately 100% of the progeny of wild type siblings of *lev-11(ETU-)/(+)* animals, or of unrelated wild type animals (Fig.3C).

Considering that previous RNAi-based studies had implicated short LEV-11 isoforms in embryonic development (Anyanful et al., 2001), we were surprised putative *lev-11(ETU-)* homozygotes did not arrest during embryogenesis, which would have been resulted in the presence of unhatched eggs (Fig.3C). To test whether the lack of a severe embryonic phenotype was due to compensation by LEV-11C, we used CRISPR/Cas9-mediated mutagenesis to insert stop codons into exon 3b, generating a *lev-11(CETU-)* allele lacking all four short LEV-11 isoforms (Fig.1). Similar to *lev-11(ETU-)*, we recovered heterozygous animals, but were unable to recover homozygotes for *lev-11(CETU-)*. Again, all eggs of *lev-11(CETU-)/(+)* heterozygotes hatched as L1 larvae, and approximately 25% of these failed to develop further (Fig.3D). Thus, larval but not embryonic development in *C. elegans* depends on zygotic expression of a short LEV-11 isoform, although these results did not rule out contributions of maternally provided short isoforms to embryogenesis.

### LEV-11E is essential for pharyngeal muscle function but not sarcomeric F-actin organization

Based on the viability of *lev-11(CDTU-)* animals and the larval arrest of *lev-11(ETU-)* and *lev-11(CETU-)* animals, we suspected LEV-11E is an essential short Tpm isoform. LEV-11E is expressed at highest levels in the excretory cell and pharyngeal muscles (Watabe et al., 2018). To test whether forced expression of LEV-11E in pharyngeal muscle is sufficient to rescue viability of *lev-11(ETU-)* and *lev-11(CETU-)* homozygotes, we cloned the *lev-11E* cDNA behind the strong *myo-2* pharyngeal muscle myosin promoter (Okkema et al., 1993) and an N-terminal mCherry tag. Plasmid DNA injected into the syncytial gonads of *C. elegans* hermaphrodites can recombine into linear extrachromosomal arrays (ECAs) that can be inherited in subsequent progeny (Mello et al., 1991). We injected plasmid bearing the mCherry-tagged LEV-11E (mCH::LEV-11E)-expressing transgene into heterozygous *lev-11(ETU-)/(+)* and heterozygous *lev-11(CETU-)/(+)* animals to produce ECAs referred to here with the shorthand notation, *Ex[mCH::lev-11E]*. Strikingly, genotyping the adult progeny of injected animals showed we could now recover homozygous *lev-11(ETU-)* and *lev-11(CETU-)* animals. Notably, these animals had mCherry-positive pharynges, indicating they had inherited the *Ex[mCH::lev-11E]* ECA. Finally, these homozygous mutants in turn produced viable mCherry-positive progeny, with no discernable loss of viability over multiple generations.

Immunoblot of whole worm extracts with anti-LEV-11(Short) confirmed *lev-11(CETU-); Ex[mCH::lev-11E]* worms lack endogenous short LEV-11 isoforms but express higher molecular weight ectopic mCH::LEV-11E (Fig.2B). Confirming this, immunostain with anti-LEV-11(Short) indicated the presence of a short LEV-11 isoform in the pharyngeal muscle of wild type animals and *lev-11(CETU-)* animals bearing *Ex[mCH::lev-11E]*, but not in *lev-11(CETU-)* animals lacking the ECA (Fig.4A). As expected, anti-LEV-11(Short) immunostain of *lev-11(CETU-); Ex[mCH::lev-11E]* animals coincided with anti-mCherry immunostain in the pharynx (Fig.4A). Interestingly, immunoblot of *lev-11(ETU-); Ex[mCH::lev-11E]* extracts also showed little to no endogenous short LEV-11 (Fig.2B), suggesting LEV-11C is likely a low abundance isoform, while LEV-11E may be the most abundant short isoform. Thus, mCherry-tagged LEV-11E can provide for all essential maternal and zygotic functions of short LEV-11 isoforms.

**Figure 4.**
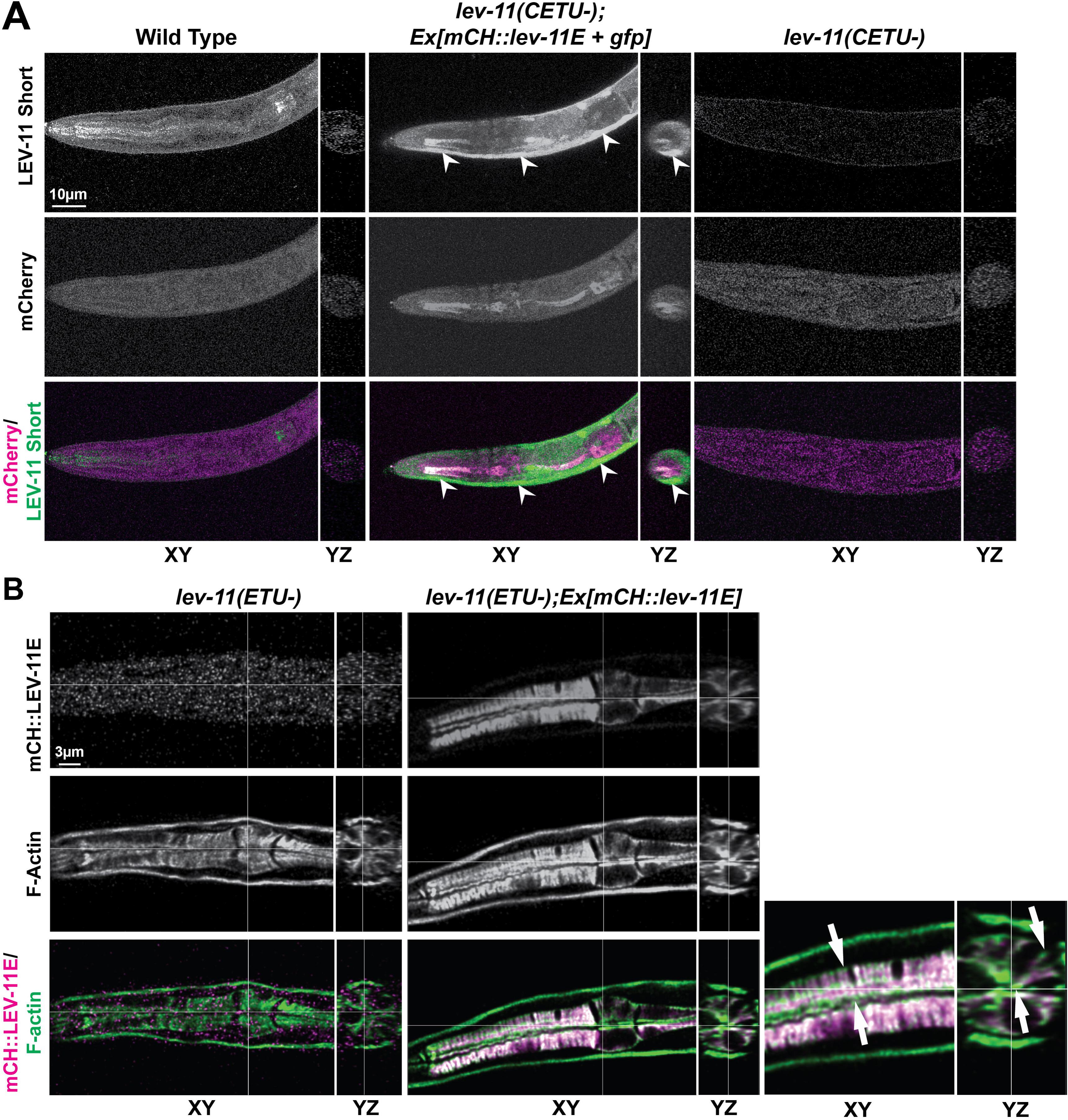
LEV-11E associates with but is not required to organize thin filaments in pharyngeal muscle sarcomeres. **(A)** Deconvolved maximum intensity projections of the XY plane show the pharynx, with YZ cross sections through pharyngeal muscle pm3, in L1 larvae immunostained for short LEV-11 isoforms and for mCherry. Anti-LEV-11(Short) similarly stains endogenous LEV-11 and mCherry-tagged LEV-11E in pharyngeal muscle, but does not stain pharyngeal muscle in *lev-11(CETU-)* worms lacking all short LEV-11 isoforms. The green channel signal observed in body wall muscle of *Ex[mCH::lev-11E]*-bearing animals (*arrowheads*) reflects the presence of a body wall muscle-expressing *GFP* transgene that is coinherited as part of the *Ex[mCH::lev-11E]* array. **(B)** Single deconvolved XY confocal plane images showing pharyngeal muscles pm3-5, with YZ cross sections through pm4, in L1 larvae stained for F-actin (*green*) and expressing or not expressing mCH::LEV-11E (*magenta)* from *Ex[mCH::lev-11E]*. Shown in enlarged XY and YZ views (*right*), mCH::LEV-11E colocalizes along F-actin structures except at the ends of filament bundles near the basal and apical muscle cell surfaces (arrows). Absence of mCH::LEV-11E in worms lacking LEV-11E, T, and U does not disturb overall F-actin organization.

Among *Ex[mCH::lev-11E]*-bearing animals, mCH::LEV-11E was strongly expressed in pharyngeal muscle, but there was also weak expression in body wall muscle, and we could not rule out unobserved lower expression in other tissues. To specifically test whether expression of LEV-11E in the pharynx is essential for viability, we utilized the propensity of ECAs to be inherited in a mosaic manner (Mello et al., 1991). We fixed L1-stage progeny of *lev-11(ETU-); Ex[mCH::lev-11E]* and of *lev-11(CETU-); Ex[mCH::lev-11E]* worms, stained these for F-actin, and inspected for mCH::LEV-11E expression in the pharyngeal muscles pm3 through pm7, ignoring pm1, pm2, and pm8 due to their relatively small sizes. For both strains, progeny at the L1 stage included animals with no pharyngeal mCH::LEV-11E, animals with mCH::LEV-11E in all pharyngeal muscle cells, and animals with mCH::LEV-11E expression in only some muscle cells (Fig.3E). When we performed a similar analysis on the progeny that had reached adulthood, > 97% of the animals expressed mCH::LEV-11E in all pharyngeal muscle cells (Fig.3E), while the remaining animals lacked mCH::LEV-11E expression in no more than one out of three muscle cells in a single pharyngeal muscle. These results suggest LEV-11E must be expressed in all the major pharyngeal muscles for normal larval development.

Pharyngeal muscle cells also serve as epithelial cells that line the pharyngeal lumen. Most pharyngeal muscle cells have large radially-arranged sarcomeres that span the distance from the apical surface of the muscle cell facing the lumen to the basal surface (Albertson et al., 1997). Thus, thin filaments attach to sarcomere Z-line equivalents at either the apical or the basal plasma membranes. In *lev-11(ETU-); Ex[mCH::lev-11E]* and *lev-11(CETU-); Ex[mCH::lev-11E]* L1 larvae and adults stained for F-actin, mCH::LEV-11E overlapped with thin filaments except near the Z-lines at the apical and basal surfaces (Fig.4B, arrows). Surprisingly, actin filament distribution was not visibly altered in *lev-11(ETU-)* or *lev-11(CETU-)* L1 animals that failed to inherit the *Ex[mCH::lev-11E]* array (Fig.4B), suggesting LEV-11E is not required to maintain thin filament stability or overall sarcomere organization. To test whether LEV-11E is required for pharyngeal muscle activity, we observed pharynges of live animals over time. In the presence of food, pharyngeal muscle of wild type animals undergoes rhythmic contraction termed “pharyngeal pumping”, which can be observed through the transparent body of worms using a stereomicroscope. We observed pharyngeal pumping in 99% of wild type L1 larvae and in 76% of mCherry-positive *lev-11(CETU-); Ex[mCH::lev-11E]*, but in only 2.7% of mCherry-negative *lev-11(CETU-)* larvae lacking LEV-11E expression (Fig.3F, Movie 1, Movie 2). Taken together, these results suggest LEV-11E plays a little to no role in organizing sarcomere structure of pharyngeal muscles, but does regulate contraction.

### FHOD-1 and LEV-11E localize and function independently in pharyngeal muscle

FHOD-1 in pharyngeal muscle concentrates at the apical and basal membranes near the ends of sarcomeric F-actin bundles (Mi-Mi et al., 2012). To determine whether this localization is affected by absence of LEV-11E, we crossed a FHOD-1::GFP-expressing transgene into the *lev-11(ETU-); Ex[mCH::lev-11E]* background. In L1 larvae expressing mCH::LEV-11E, FHOD-1::GFP localized at the ends of mCH::LEV-11E-decorated filament bundles near the Z-lines at the plasma membranes, but with little overlap with LEV-11E (Fig.5, arrows). This suggests FHOD-1 and LEV-11E bind distinct domains on pharyngeal thin filaments. In *lev-11(ETU-)* animals that failed to inherit the mCH::LEV-11E-expressing ECA, pharyngeal FHOD-1::GFP localization was not noticeably changed, suggesting FHOD-1 localization does not depend on LEV-11E (Fig.5).

**Figure 5.**
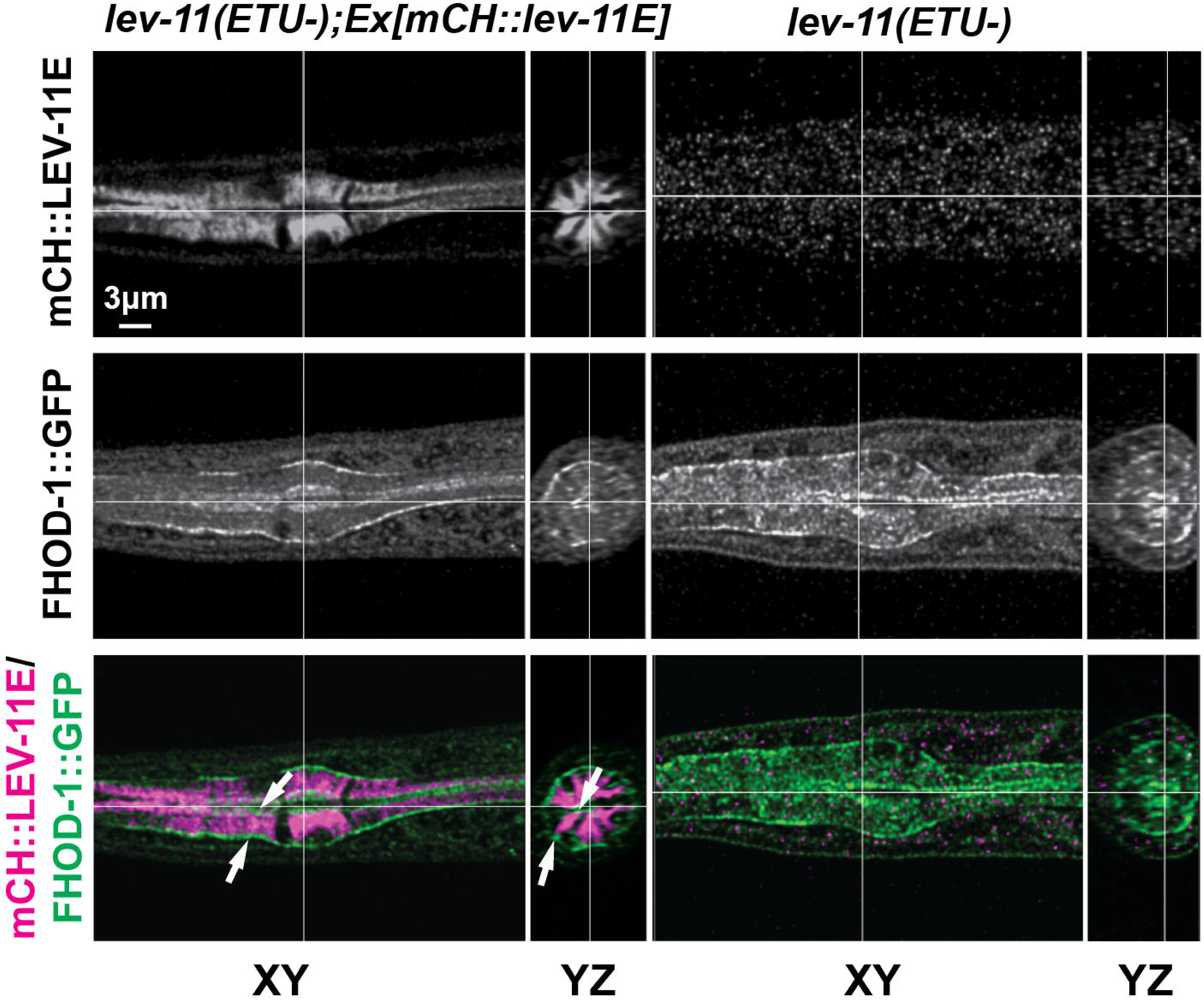
FHOD-1::GFP localizes near the barbed ends of thin filaments in pharyngeal muscle independently of LEV-11E. Single deconvolved XY confocal plane images showing pharyngeal muscles pm3-5, with YZ cross sections through pm4, in L1 larvae show FHOD-1::GFP is concentrated at the apical and basal surfaces of pharyngeal muscle cells, particularly at the ends of mCH::LEV-11E-positive filament bundles (*arrows*). Localization of FHOD-1::GFP is not significantly changed in *lev-11(ETU-)* mutants that express no LEV-11E.

To test whether pharyngeal LEV-11E depends on FHOD-1 for localization, we crossed *fhod-1(*Δ) into the *lev-11(CETU-); Ex[mCH::lev-11E]* genetic background. As we have seen previously *for fhod-1(*Δ) (Mi-Mi et al., 2012), pharyngeal F-actin appeared normal in *fhod-1(*Δ*) lev-11(CETU-); Ex[mCH::lev-11E]* L1 larvae and adults expressing mCH::LEV-11E (Fig.6A.B). Moreover, mCH::LEV-11E localized normally in *fhod-1(*Δ) animals (Fig.6A,B), and such animals exhibited pharyngeal pumping, were viable, and grew to adulthood. Thus, FHOD-1 does not appear to influence the function or localization of LEV-11E.

**Figure 6.**
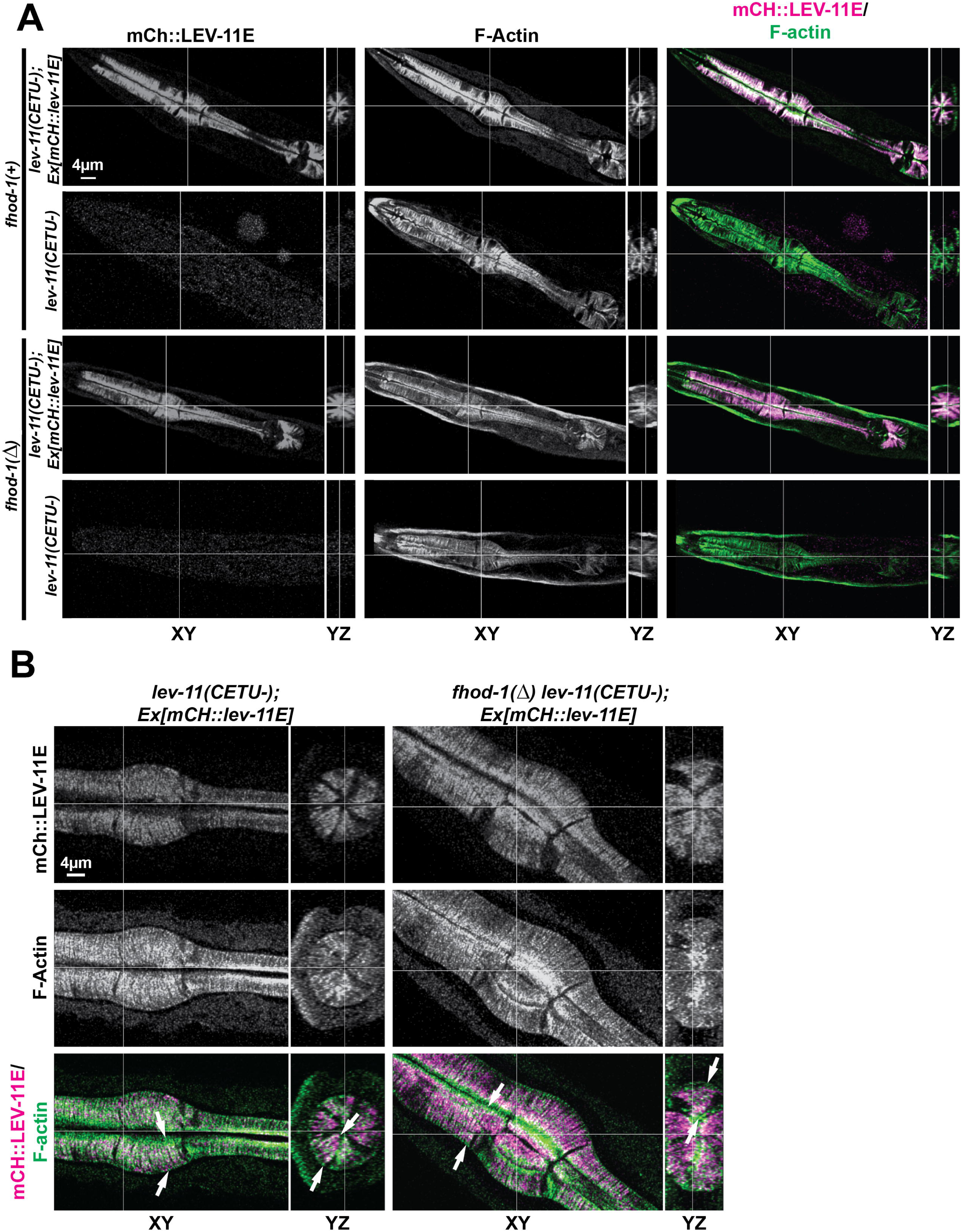
Absence of FHOD-1 and short LEV-11 isoforms do not substantially disturb actin organization in pharyngeal muscle. **(A)** Single deconvolved XY confocal slices showing the pharynx (all pms), with YZ cross sections through pm4, in L1 larvae stained for F-actin. Thin filament organization is not significantly altered in pharyngeal muscle in animals lacking FHOD-1 or short LEV-11 isoforms. Additionally, mCH::LEV-11E organization is not significantly altered in the pharyngeal muscle of *fhod-1(*Δ) larvae. **(B)** Single deconvolved XY confocal slices showing pm3-5, with a YZ cross sections through pm4, in day 1 adults stained for F-actin. Even in absence of FHOD-1 in *fhod-1(*Δ) animals, mCH::LEV-11E localizes along sarcomeric thin filaments except immediately adjacent to sarcomere Z-lines at the apical and basal membranes (*arrows*).

To determine whether simultaneous loss of FHOD-1 and LEV-11E has a synthetic effect on pharyngeal sarcomere organization, we examined F-actin in *fhod-1(*Δ*) lev-11(CETU-)* L1 larvae that failed to inherit the mCH::LEV-11E-expressing ECA. Again, pharyngeal F-actin appeared normal in these animals (Fig.6A). Thus, FHOD-1 and LEV-11E associate with different domains of sarcomeric thin filaments in pharyngeal muscle, neither protein affects the localization of the other, and both proteins are dispensable to establishing the overall actin cytoarchitecture of pharyngeal muscle.

### Short isoforms LEV-11C and E can bind thin filaments in body wall muscle when exogenously expressed, but are not important for FHOD-1 functions in body wall muscle

Short LEV-11 isoforms co-precipitated with FHOD-1 from extracts enriched for body wall muscle sarcomere proteins. Thus, we also probed for short LEV-11 isoform expression and function in body wall muscle. FHOD-1 in body wall muscle localizes to small bodies at the edges of muscle cells and as faint striations that overlap dense bodies, the primary sarcomere Z-line structures (Mi-Mi et al., 2012). Long LEV-11 isoforms are highly expressed in body wall muscle (Barnes et al., 2018; Watabe et al., 2018), and previous immunostain with a pan-LEV-11 antibody has shown LEV-11 decorates thin filaments except for a segment proximal to the dense bodies/Z-lines (Ono & Ono, 2002; Barnes et al., 2018). GFP-tagged short isoform LEV-11T (GFP::LEV-11T) exogenously expressed in body wall muscle adopts a similar distribution on thin filaments, while LEV-11U, which a much lower affinity for F-actin, forms aggregates when overexpressed in body wall muscle (Ono et al., 2023). We generated ECAs expressing GFP-tagged short isoforms LEV-11C or E, or long isoform LEV-11A using the strong body wall muscle-specific *myo-3* promoter (Waterston, 1989) in worms with endogenous dense body marker α-actinin/ATN-1 tagged with mCherry (ATN-1::mCH) (Kimmich et al., 2024). As expected, GFP::LEV-11A colocalized with thin filament F-actin except in a zone proximal to the dense bodies (Fig.7A). Similar to LEV-11T, we also observed this localization for GFP::LEV-11C and GFP::LEV-11E (Fig.7A), suggesting multiple short isoforms of LEV-11 have the capacity to bind thin filaments in body wall muscle sarcomeres.

**Figure 7.**
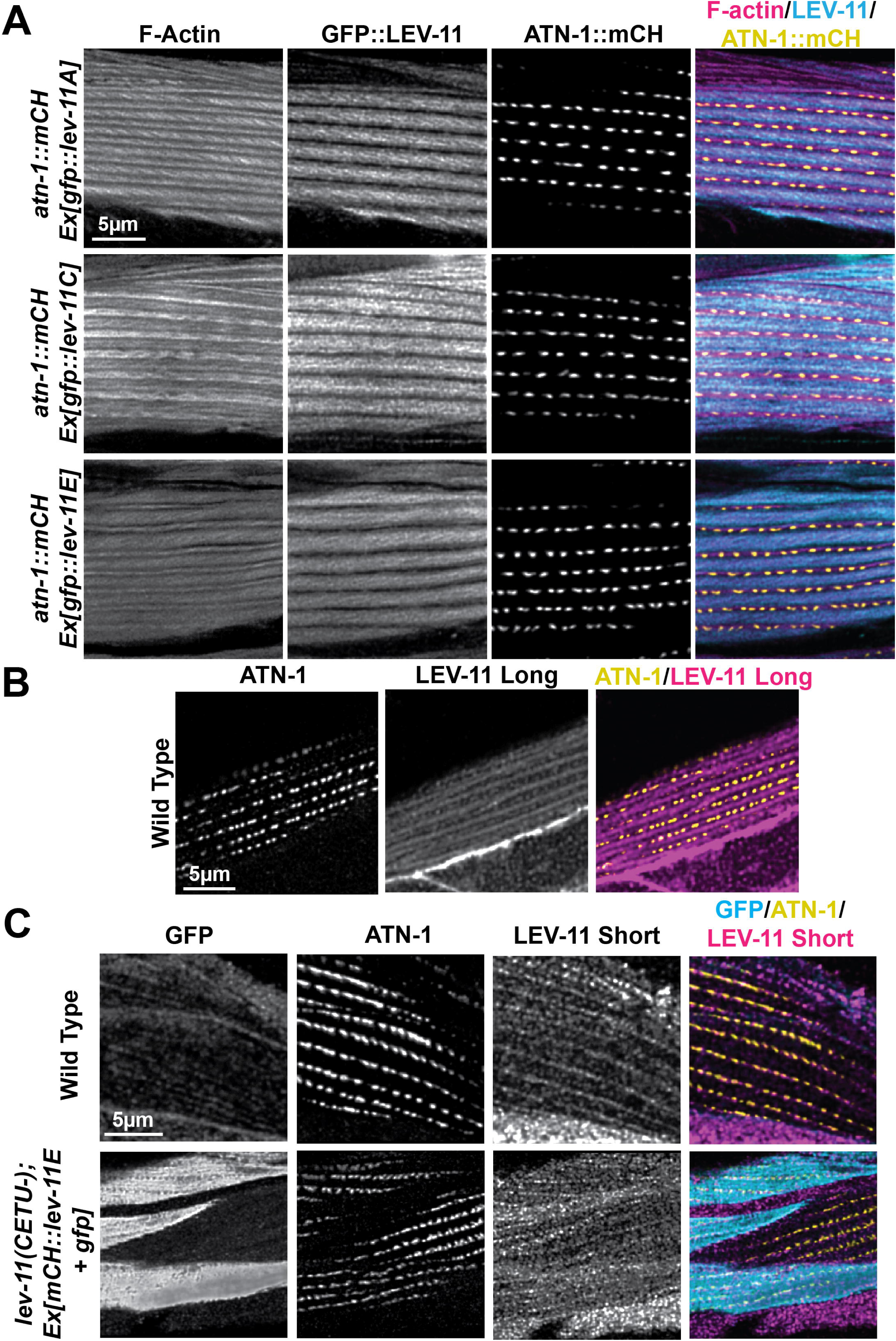
Short LEV-11 isoforms bind thin filaments in body wall muscle when ectopically expressed, but are not endogenously present in body wall muscle sarcomeres. **(A)** Single deconvolved confocal slices of body wall muscle of day-1 adults expressing mCherry-tagged ATN-1 and the indicated GFP-tagged LEV-11 isoforms were stained for F-actin. All three LEV-11 isoforms localize to thin filaments based on strong overlapping signal with F-actin, but are not present immediately adjacent to mCH::ATN-1-positive dense bodies. **(B)** Single deconvolved confocal slices of body wall muscle of a wild type day-1 adult immunostained with anti-LEV-11(Long) and anti-ATN-1 shows a striated distribution of LEV-11 flanking ATN-1-positive dense bodies. **(C)** Single deconvolved confocal slices of day-1 adult worms immunostained for ATN-1, GFP (a co-expression marker for the mCH::LEV-11E-expressing array), and short LEV-11 isoforms shows anti-LEV-11(Short) immunostain along striations that intersect dense bodies in wild type animals and in *lev-11(CETU-)* mutants, regardless of whether body wall muscle cells inherited the mCH::LEV-11E + GFP-expressing array.

To probe for endogenous LEV-11 expression in body wall muscle, we immunostained wild type animals using anti-LEV-11(Long) or anti-LEV-11(Short), and an antibody that recognizes the dense body protein ATN-1 (Fig.7B). As expected, anti-LEV-11(Long) immunostain recapitulated our observations with GFP::LEV-11A (Fig.7B) and previous pan anti-LEV-11 immunostain (Ono & Ono, 2002). However, immunostain with anti-LEV-11(Short) did not resemble over-expressed GFP-tagged short isoforms, but instead decorated faint striations in body wall muscle that overlapped dense bodies (Fig.7C). To test the specificity of this immunostain, we also stained anti-LEV-11(Short) in *lev-11(CETU-)* animals bearing a modified ECA, *Ex[mCH::lev-11E + myo::gfp]*. This ECA expresses mCH::LEV-11E from the pharyngeal *myo-2* promoter, but at a lower level than previously to reduce the leaky body wall muscle expression. Additionally, the ECA carries a GFP transgene driven by the *myo-3* promoter for strong body wall muscle expression, allowing the presence of GFP to indicate inheritance of the ECA by individual body wall muscle cells. Thus, GFP-negative body wall muscle cells would lack the mCH::LEV-11 transgene (and therefore would lack all short LEV-11 isoforms). Immunostain anti-LEV-11(Short) produced a striated pattern identical to that seen in wild type animals in all body wall muscle cells of this strain, regardless of whether cells were GFP-positive or GFP-negative (Fig.7C). These results strongly suggest anti-LEV-11(Short) immunostain in body wall muscle is non-specific.

Despite our failure to detect endogenous short LEV-11 isoforms in body wall muscle sarcomeres, we tested whether they might contribute to FHOD-1 functions in body wall muscle. FHOD-1 promotes new sarcomere formation in body wall muscle, and in cooperation with profilin, promotes the stability of dense bodies (Mi-Mi et al., 2012; Mi-Mi & Pruyne, 2015; Sundaramurthy et al., 2020; Kimmich et al., 2024). FHOD-1 promotes new sarcomere formation in body wall muscles, such that body wall muscle cells in *fhod-1(*Δ) mutants assemble striations more slowly than in wild type animals, resulting in narrow body wall muscles with fewer striations per muscle cell (Mi-Mi et al., 2012; Sundaramurthy et al., 2020; Kimmich et al., 2024). Examining body wall muscle size in the three *lev-11(C-)* strains, one strain had narrower body wall muscle muscles similar to, but of lesser magnitude than, *fhod-1(*Δ) mutants, as compared to age-matched wild type animals (Fig.8A). But no such difference was observed for the other two strains, suggesting this effect was unrelated to *lev-11(C-)*. Additionally, no *lev-11(C-)* strains had significantly fewer striations per muscle cell than wild type (Fig.8B), unlike *fhod-1(*Δ) mutants. Thus, LEV-11C is not required for FHOD-1-dependent body wall muscle growth. We chose not to make a similar comparison for mutants lacking LEV-11E, as partial defects in pharyngeal pumping and feeding can lead to reduced body growth, which negatively impacts body wall muscle growth (Sundaramurthy et al., 2020). However, FHOD-1 performs an additional function in body wall muscle, working in cooperation with profilin to promote stability in dense bodies in body wall muscle (Kimmich et al., 2024). Whereas dense bodies in wild type animals appear relatively uniform in size, shape, and spacing, those in *fhod-1(*Δ) mutants appear irregular in shape and spacing (Sundaramurthy et al., 2020; Kimmich et al., 2024). We immunostained for the dense body protein ATN-1 and the ECA marker GFP in *lev-11(CETU-)* animals with mosaic inheritance of *Ex[mCH::lev-11E + myo::gfp]*. Dense bodies appeared normal in the body wall muscle cells of these animals regardless of the presence or absence of the ECA (Fig.8C), suggesting absence of all short LEV-11 isoforms does not affect dense bodies in a manner similar to *fhod-1(*Δ). In summary, we did not detect endogenous short LEV-11 isoforms in body wall muscle sarcomeres, or find evidence short LEV-11 isoforms function with FHOD-1 in body wall muscle development.

**Figure 8.**
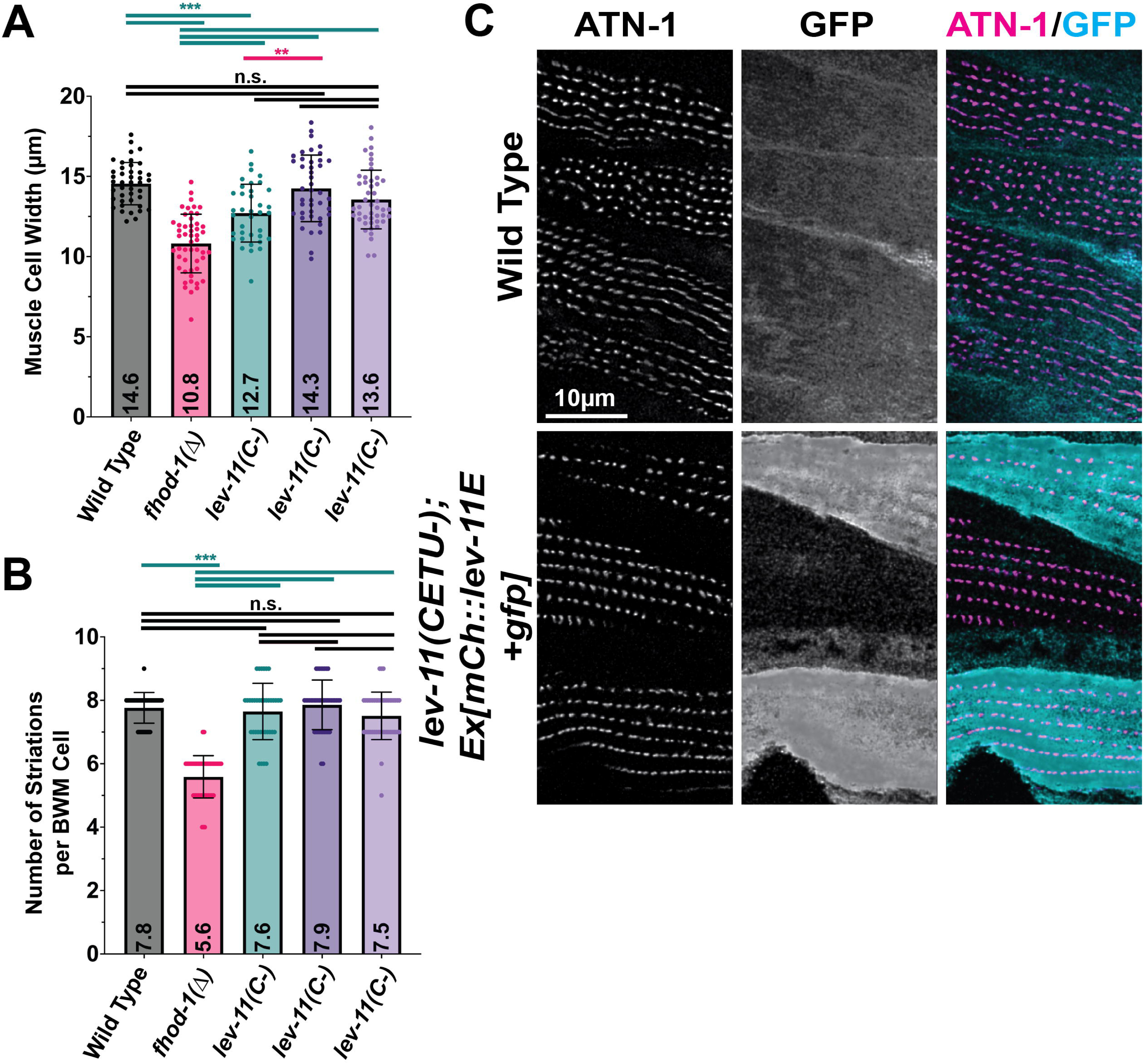
Mutations disrupting short LEV-11 isoforms do not result in body wall muscle defects. **(A, B)** Adult worms were stained for F-actin and **(A)** body wall muscle cell widths were measured, and **(B)** the number of striations per body wall muscle cell were counted (n = 15-20 animals per strain, 2 - 4 muscle cells per animal). (**) *p* = 0.0016; (***) *p* < 0.0001; (n.s.) not significant, *p* ≥ 0.05. **(C)** Single deconvolved confocal slices of body wall muscle in worms immunostained for dense body marker ATN-1 and for GFP, a marker for the presence of an ECA bearing an mCH::LEV-11E-expressing transgene. Dense bodies have relatively uniform appearance and spacing along striations in both strains, including in GFP-negative cells of *lev-11(CETU-)* animals that lack all short LEV-11 isoform expression.

## DISCUSSION

*C. elegans* provides an ideal model to systematically analyze the core functions of Tpms in an animal, having a relatively simple anatomy and a single Tpm-coding gene (*lev-11*) that produces a smaller complement of Tpm splice isoforms than in vertebrates. However, similar to vertebrate Tpms, the worm splice isoforms are a mix of higher molecular weight proteins (long isoforms) generated from an early promoter of the *lev-11* gene, and lower molecular weight proteins (short isoforms) generated from an internal promoter. Through a combination of promoter analysis and fluorescent reporters for alternative splicing, the primary tissue distributions for five of the seven known splice isoforms of LEV-11 have been determined (Kagawa et al., 1995; Anyanful et al., 2001; Barnes et al., 2018; Watabe et al., 2018). Three long isoforms are expressed primarily in the striated body wall muscles, with one splice form (LEV-11O) expressed in the anterior-most muscle cells, and the others (LEV-11A and D) expressed in the remaining body wall muscle, where they regulate actomyosin activity in their respective muscle cells (Barnes et al., 2018). Of the short Tpm isoforms, LEV-11C is primarily expressed in the intestine and neurons, and LEV-11E is expressed in pharyngeal muscle and the excretory cell, while short isoforms LEV-11T and LEV-11U appear to be low abundance proteins, with unknown tissue distributions (Watabe et al., 2018; Ono et al., 2023). Short Tpm isoforms have been shown to play a wide variety of roles in cells, ranging from regulating cell migration through controlling proliferation (Bach et al., 2009; Lees et al., 2013; Cagigas et al., 2022). Here, we used a CRISPR-Cas9-based mutagenesis strategy to determine the minimal required complement of short Tpm isoforms in an intact animal, and remarkably determined *C. elegans* can survive with a single short isoform, LEV-11E.

To abolish all short Tpm isoforms while having no effect on the coding sequences of any long isoforms, we introduced premature stop codons into the starting exon shared by all short isoforms. Western blot confirmed this *lev-11(CETU-)* allele abolished endogenous short Tpms (Fig.2B). Worms heterozygous for *lev-11(CETU-)* are grossly normal, but their homozygous *lev-11(CETU-)* progeny arrest and die during very early larval development (Fig.3D), demonstrating a requirement for zygotic expression of one or more short Tpm isoforms for progression beyond the L1 larval stage. Notably, we did not observe any embryonic lethality among the progeny of animals heterozygous for *lev-11(CETU-)* (Fig.3D). This result differed from a previous RNAi-based study that demonstrated animals inhibited for short Tpm isoform expression in the germline generate progeny with frequent failures in embryogenesis and larval development (Anyanful et al., 2001). A possible explanation for this discrepancy is that genetic loss-of-function mutants we generated would still receive a maternal provision of short Tpm mRNA and/or protein from their heterozygous parent, which could sustain them through embryogenesis. In contrast, RNAi treatment of the germline would inhibit both maternal and zygotic expression of short Tpm in the progeny of treated animals. Thus, our results do not address possible roles for short Tpm isoforms in embryogenesis, or in the earlier processes of gametogenesis.

Through mutation of additional *lev-11* exons, we showed the larval arrest of *lev-11(CETU-)* animals is associated with the elimination of LEV-11E but not the other isoforms. Thus, animals homozygous for disruption of an exon used uniquely by LEV-11C, or for an exon shared by the short isoforms LEV-11C, T, and U, were viable and grossly normal appearing. In contrast, homozygous disruption of an exon shared by LEV-11E, T, and U resulted in early larval arrest identical to that of *lev-11(CETU-)* animals (Fig.3A,C). Moreover, re-expression of a LEV-11E-coding transgene restored viability to *lev-11(CETU-)* mutants. Again, these results differ somewhat from Anyanful and colleagues (2001), who observed RNAi-based targeting of the LEV-11C-specific exon resulted in embryonic and larval defects similar to that for RNAi targeting all short isoforms (Anyanful et al., 2001). We suspect this discrepancy may be due to transitive RNAi, in which RNA-dependent RNA polymerases in *C. elegans* generate additional small interfering RNAs (siRNAs) from sequences 5’ to the originally targeted mRNA region, spreading an RNAi effect (Sijen et al., 2001; Alder et al., 2003). In this case, RNAi targeting the LEV-11C-specific exon 5c might have resulted in transitive RNAi targeting exons 3b and 4a, which are shared by all short Tpm splice isoforms in *C. elegans*. Our results do not rule out important roles for short isoforms LEV-11C, T, and U, as we only examined gross morphology and fertility. Indeed, a previous study has shown animals disrupted for LEV-11U have modest movement and brood size defects (Ono et al., 2023), while strong expression of LEV-11C in the intestine and neurons suggests a role for that isoform in those tissues (Watabe et al., 2018).

Among vertebrates, many long Tpm isoforms associate with thin filaments in muscle sarcomeres, and are thus called “muscle” or “sarcomeric” Tpms. In contrast, vertebrate short Tpm isoforms (as well as some long isoforms) are often termed “nonmuscle” or “nonsarcomeric” Tpms because they are abundantly expressed in nonmuscle cell types, or are expressed in muscle cells but in association with nonsarcomeric actin filaments. For example, mouse short Tpm isoforms Tm5NM1 and Tm4 associate with actin filaments positioned outside sarcomeres but adjacent to Z-disks, and Tm4 associates with extra-sarcomeric filaments involved in muscle repair (Kee et al., 2004; Vlahovich et al., 2007; Vlahovich et al., 2009). By these criteria, the short isoform LEV-11C expressed in the intestine and in neurons would be considered a nonmuscle Tpm (Watabe et al, 2018). However, we detected one or more short LEV-11 isoforms as co-precipitants during immunoprecipitations of FHOD-1, a formin primarily expressed in muscle in the worm (Table 1; Mi-Mi et al., 2012). We and others showed that when GFP-tagged short isoforms LEV-11C, LEV-11E, and LEV-11T are overexpressed in body wall muscle, they associate with sarcomeric thin filaments (Fig.7A; Ono et al., 2023), but we found no evidence of endogenous expression of any short LEV-11 isoform in body wall muscle (Fig.7C). In contrast, in pharyngeal muscle, LEV-11E is the only Tpm that is abundantly expressed (Watabe et al., 2018), and our results suggest it functions there as a sarcomeric Tpm. Endogenous LEV-11E and exogenous mCherry-tagged LEV-11E associate with thin filaments of pharyngeal muscle sarcomeres (Fig.4). Analysis of mosaic inheritance of a *lev-11E* transgene in *lev-11(CETU-)* animals showed development beyond the early L1 larval stage depends on expression of LEV-11E in all major pharyngeal muscle cells (Fig.3E), and animals that lack pharyngeal LEV-11E exhibit pharyngeal muscle paralysis (Fig.3F), which would lead to starvation and arrested development. We were not able to observe expression of the *lev-11E* transgene in tissues other than pharyngeal and body wall muscle, and thus we are unable to address whether LEV-11E performs additional essential functions in other tissues, such as the excretory cell where LEV-11E is also highly expressed (Watabe et al., 2018).

In different contexts, Tpms have been shown to affect sarcomere structure and function. Striated muscles host actin filament severing proteins such as gelsolin and ADF/cofilin (Rouayrenc et al., 1984; Ono et al., 1993, 1993, 1994, 1999), yet thin filaments tend to be stable structures with relatively slow turnover (Skwarek-Maruszewska et al., 2009), suggesting they are stabilized against filament disassembly. Tpms in general inhibit severing by these proteins (Gunning et al., 2015), and worm LEV-11 potently antagonizes filament severing by the body wall muscle ADF/cofilin homolog UNC-60B (Ono & Ono, 2002). However, F-actin organization appears relatively normal in LEV-11E-deficient pharyngeal muscle cells (Figs 4-6), suggesting that thin filament stability is not wholly dependent on LEV-11E. This differs from what has been observed in body wall muscle, where partial knockdown of either all isoforms or all long isoforms of LEV-11 partially disrupt actin organization in body wall muscle in an ADF/cofilin-dependent manner (Ono & Ono, 2002). Tpms also regulate actomyosin contractility in muscle by controlling the degree to which myosin accesses binding sites on thin filaments (Geeves, 2012). Based on the failure of LEV-11E-deficient pharyngeal muscles to rhythmically pump, we suggest LEV-11E similarly controls actomyosin activity in pharyngeal muscle sarcomeres.

Based on the strong expression of FHOD-1 and LEV-11E in pharyngeal muscle, and the apparent absence of endogenous short LEV-11 isoforms in body wall muscle, we suggest our co-precipitation of FHOD-1 with short LEV-11 isoforms (Table 1) reflected capture of pharyngeal muscle thin filaments. Immunostain suggests FHOD-1 and LEV-11E decorate the same sarcomeric thin filaments in pharyngeal muscle, but on different locations. That is, FHOD-1 is present near the sarcomere Z-lines at the plasma membrane, while LEV-11E decorates the remaining portions of the thin filaments distal to this location (Fig.5). Interestingly, this resembles the spatial relationship between FHOD-1 and long LEV-11 isoforms in body wall muscle, where FHOD-1 is present very near dense bodies that serve as Z-lines, while long LEV-11 isoforms occupy the portions of thin filaments more distant from the dense bodies (Fig.7; Ono & Ono, 2002).

Formins and Tpms have been shown to functionally interact in several contexts. For example, Tpms have been shown to enhance formin-mediated actin polymerization *in vitro* for several formins, while different formin isoforms in fission yeast direct preferential assembly of different Tpm isoforms onto actin filaments *in vivo* (Wawro et al., 2007; Johnson et al., 2014; Alioto et al., 2016). However, we found no direct phenotypic evidence for such formin/Tpm cooperation between FHOD-1 and LEV-11E. We observed no effect of loss of FHOD-1 on pharyngeal muscle contractility, F-actin organization, or LEV-11E distribution (Fig.6), and in turn, FHOD-1 does not depend on LEV-11E for its localization to pharyngeal muscle thin filaments (Fig.5). Even simultaneous absence of FHOD-1 and LEV-11E has no noticeable effect on pharyngeal muscle actin organization (Fig.6A). In body wall muscle, where FHOD-1 is required for normal dense body morphogenesis and stability (Sundaramurthy et al., 2020; Kimmich et al., 2024), short Tpm isoforms do not appear to play a role (Fig.8C).

In summary, we find that the simple animal model *C. elegans* requires only a single short Tpm isoform for development and fertility, and in contrast to short Tpm isoform functions in vertebrates, at least one essential role for this short isoform is as a sarcomeric Tpm in pharyngeal muscle.

## MATERIALS AND METHODS

### Plasmids

*lev-11E* cDNA was isolated from Yuji Kohara clone yk1035c01 with a single point mutation (T667A coding V225A) that we corrected by site-directed mutagenesis using QuikChange Lightning (Agilent, Santa Clara, CA). *lev-11C* cDNA was isolated from Yuji Kohara clone yk1213f01. *lev-11A* cDNA was isolated from the Pro-Quest *C. elegans* cDNA library for yeast two hybrid screening (discontinued product from Invitrogen). Plasmids L4440-lev-11C and L4440-lev-11E for dsRNA production for RNAi were generated by amplifying the first three exons of *lev-11C* and *lev-*11E, respectively, with flanking *KpnI* and *NheI* sites, and subcloning into L4440. Plasmid pLMM19 was generated by introducing flanking *BamHI* sites to *lev-11C* cDNA by PCR and subcloning into pGEX-6p-3. Plasmid pLMM25 was generated by introducing flanking *SmaI* sites to *lev-11E* cDNA and subcloning into pGEX-6p-2. Plasmid pGEX-6p-3-lev-11A was generated by seamlessly cloning *lev-11A* cDNA into pGEX-6p-3 by In-Fusion cloning (Takara, Shiga, Japan). For expression of mCherry-tagged LEV-11 isoforms in body-wall muscle, *lev-11A*, *C*, and *E* cDNAs were appended at their 5’ ends with sequence encoding 5-amino acid Ala-Gly-Ala-Gly-Ser linker, and flanked by *BamHI* sites using PCR before insertion by In-Fusion cloning into pCFJ104 (Addgene #19328; Watertown, MA), between the body wall muscle-specific *myo-3* promoter driving mCherry, and the *unc-54* 3’ untranslated region (UTR). Resultant plasmids were named pCP-Cherry-lev-11A, pCP-Cherry-lev-11C, and pCP-Cherry-lev-11E. For pharyngeal expression of LEV-11E, *lev-11E* cDNA from pCP-Cherry-lev-11E, together with linker and *BamHI* sites, was amplified and inserted by In-Fusion cloning between *myo-2* promoter driving mCherry and *unc-54* 3’UTR of pCFJ90 (Addgene #19327; Watertown, MA) to generate plasmid pKM3. Plasmid *myo-3::gfp* was created by fusion PCR of the *myo-3* promoter, *gfp* (from pPD95.75), and the *unc-54* 3’ UTR, followed by TOPO-TA cloning of the resultant *myo-3p::gfp::unc-54 3’utr* product into plasmid pCR2.1 (Thermo Fisher Scientific, Waltham, MA). Plasmids *myo-2::gfp* and *rab-3::gfp* were created in a similar manner, using the *myo-2* or *rab-3* promoters, respectively, in place of the *myo-3* promoter. For expression of GFP-tagged LEV-11 isoforms in body-wall muscle, *lev-11A*, *C*, and *E* cDNAs each appended at the 5’ end with a Gly-Ala-Gly-Ala-Ser-coding linker sequence and flanked by *NheI* sites using PCR, and inserted into plasmid *myo-3::gfp* by In-Fusion cloning, generating plasmids pCP-GFP-lev-11A, pCP-GFP-lev-11C, and pCP-GFP-lev-11E, respectively. All construct sequences were verified through Sanger sequencing (Genewiz, South Plainfield, NJ).

### Worm strains and growth conditions

Worms were maintained under standard growth conditions at 20°C on nematode growth medium (NGM) plates with OP50-1 *Escherichia coli* food (Brenner, 1974). Age synchronization was done by dissolving gravid adults in 1:2 ratio reagent grade bleach:5 M NaOH to liberate embryos, and hatching embryos overnight at 20°C in M9 buffer (Ausubel et al., 1992).

Worms were grown in liquid culture for IP-MS experiments. Briefly, ten adult worms were bleached to release embryos, which were grown at 20°C on NGM plates with OP50-1 until they had reached adulthood. Twenty adult worms were then moved to five new plates and grown at 20°C. After 84 hr, worms were washed off plates with sterile M9 buffer and transferred into a 2.8 L beveled flask containing OP50-1 bacteria suspended in 500 mL of S-Complete medium (Sulston & Hodgkin, 1988), and grown at 20°C while shaking (∼ 200 rpm). When the majority of worms in the liquid culture had become adults, sucrose gradient centrifugation was applied to the culture to separate the live adult worms from the debris. Worms were washed with ice-cold sterile M9 buffer then 0.1 M NaCl solution, and then treated with alkaline bleach solution for 5-7 min to release embryos. Embryos were washed with ice-cold sterile water and allowed to hatch in M9 buffer overnight at 20°C. The next day, all newly hatched and starved L1-stage worms were transferred into a 2.8 L Flask containing OP50-1 bacteria in 500 mL of S-Complete medium and allowed to grow as an age-synchronized culture at 20°C until they reached the desired developmental stage.

Worms were treated for RNAi against short LEV-11 isoforms (Hegsted et al., 2019). Briefly, wild type worms were grown for three generations on IPTG-induced HT115 *E. coli* (Wang and Barr, 2005) expressing dsRNA from plasmids L4440-lev-11C and L4440-lev-11E.

The full genotypes of all worm strains used in this study are listed in Supplementary Table S4. Strains N2 [*wild type*], CB1467 [*him-5(e1461)*] that has a high incidence of male progeny, and KR2151 [*hIn1[unc-54(h1040)]*] bearing a chromosomal inversion on chromosome I, were obtained from the Caenorhabditis elegans Genetics Center (University of Minnesota, Twin Cities, MN). Worm strains XA8001 [*fhod-1(tm2363)*] bearing a putative null allele of *fhod-1* (Mi-Mi et al., 2012), DWP3 [*qaIs8001[fhod-1::gfp]*] bearing a GFP-tagged *fhod-1* transgene (Mi-Mi et al., 2012), and RSL62 [*atn-1(ftw35 atn-1::mCherry::ICR::GFPnls)*] tagged with mCherry at the endogenous *atn-1* locus (Kimmich et al., 2024) were previously isolated. For production of male animals bearing *fhod-1(tm2363)* for genetic crosses, XA8001 was crossed with CB1467 to generate XA8032 [*fhod-1(tm2363); him-5(e1461)*].

Strategies for our first two CRISPR-Cas9-mediated mutagenesis targets, *lev-11* exons 5a and 5c, were developed and conducted in collaboration with InVivo Biosystems (Eugene, OR), resulting in alleles *lev-11(ups174)*, called *lev-11(C-)* in the text, which introduces a three-frame stop and novel *XhoI* site into exon 5c (targeting LEV-11C), and *lev-11(ups175)/lev-11(ETU-)*, which introduces a three-frame stop into and eliminates an endogenous *SalI* site from exon 5a (targeting LEV-11E, T, and U). Worm strains DWP 262, DWP 263, and DWP 264 are three independent isolates of *lev-11(ups174)*, while strains DWP 265 and DWP 266 are two independent isolates that are heterozygous for *lev-11(ups175)*. Crossing of DWP 265 [*lev-11(ups175)/(+)*] with KR2151 produced the stably balanced heterozygous strain DWP267 [*lev-11(ups175)/hIn1*]. Additional CRISPR-Cas9 mutations targeting *lev-11* were generated based on previously published protocols (Smith et al., 2020). For this, Cas9 protein (Integrated DNA Technologies, IDT, Coralville, IA), tracrRNA (IDT), crRNAs (IDT) and ssODNs (IDT) were injected into the syncytial gonad for homology-directed repair. Allele *lev-11(ups182)/lev-11(CDTU-)* was generated by inserting a 3-frame stop and silently adding a *XhoI* site into exon 9b (targeting short isoforms LEV-11C, T, and U, and long isoform LEV-11D), using crRNA [UAU UUC AGA ACU CCG UGA CGC GG] and ssODN [TAT AAA TAA CAG TGT GAC TCA ATA TTT CAG AAC TCT AAA TAA ATA AAC TCG AGA AAG CAC GCC AAT TAC AAG ATG AGC TTG ACC ATA T]. Strains DWP276, DWP277, and DWP278 are three independent isolates of *lev-11(ups182)*. Allele *lev-11(ups184)/lev-11(CETU-)* was generated by inserting a 3-frame stop and silently adding a *BamHI* site into exon 3b (targeting all four short isoforms LEV-11C, E, T, and U) using crRNA [AAU GUC GAA GGU AAA CAA GGA GG] and ssODN [ACA CCT CTG ACT TCT GCA GAA TGT CGA AGG TAA ACT AAA TAA ATA AAG GGA TCC ACA TCG CTT CTT GAT GTC CTC AAG AAG AAG ATG CG]. Two independent isolates of *lev-11(184)* were crossed with KR2151 to produce the stably balanced heterozygous strains DWP280 and DWP281 [*lev-11(ups184)*/*hIn1*]. The sequences of all CRISPR-generated *lev-11* alleles were verified by Sanger sequencing.

To express fluorescently tagged LEV-11 proteins, adult hermaphrodites were injected in the syncytial gonads with plasmids for expression of GFP- or mCherry-tagged LEV-11 isoforms and for expression of co-injection markers for visual identification of transgenic progeny or individual muscle cells. For over expression of GFP-tagged LEV-11 isoforms used to validate antibodies for western blot analysis (Fig.2A), N2 worms were injected with 1 ng/µl pCP-GFP-lev-11A, C, or E, together with 90 ng/µl pRS315 (Sikorski & Hieter, 1989) carrier DNA, and 10 ng/µl pGH8 (neuronal mCherry expression), 5 ng/µl pCFJ104 (body wall muscle mCherry expression), and 2.5 ng/µl pCFJ90 (pharyngeal mCherry expression) co-injection markers. Resulting strains DWP112 [*upsEx62[myo-3p::gfp::lev-11E]*], DWP115 [*upsEx65[myo-3p::gfp::lev-11A]*], and DWP120 [*upsEx70[myo-3p::gfp::lev-11C]*] express high levels of GFP-tagged LEV-11E, LEV-11A, and LEV-11C, respectively, in body wall muscle. To compare the distribution of GFP-tagged LEV-11 isoforms to dense bodies in body wall muscle (Fig.8A), RSL62 worms were injected with 1 ng/µl pCP-GFP-lev-11A, C, or E, together with 90 ng/µl pRS315, 10 ng/µl *rab-3::gfp*, and 2.5 ng/µl *myo-2::gfp*, generating strains DWP243 [*atn-1(ftw35[atn-1::mCH]); upsEx161[myo-3p::mCH::lev-11A]*], DWP250 [*atn-1(ftw35[atn-1::mCH]); upsEx168[myo-3p::mCH::lev-11C]*], and DWP251 [*atn-1(ftw35[atn-1::mCH]); upsEX169[myo-3p::mCH::lev-11E]*], respectively. To drive pharyngeal expression of mCherry-tagged LEV-11E in homozygous *lev-11(ups175)* animals, strain DWP267 was injected with 2.5 ng/µl pKM3, 90 ng/µl pRS315, and 10 ng/µl *rab-3::gfp* plasmid, resulting in a viable homozygous *lev-11(ups175)* strain DWP275 [*lev-11(ups175); upsEx178[myo-2p::mCH::lev-11E]*]. Similarly, DWP280 [*lev-11(ups184)/hIn1*] was injected with 2.5 ng/µl pKM3, 37.5 ng/µl pRS315, 10 ng/µl *rab-3::gfp* plasmid, and 50 ng/µl *myo-3::gfp* plasmid, resulting in a viable homozygous *lev-11(ups184)* strain, DWP282 [*lev-11(ups184); upsEx185[myo-2p::mCH::lev-11E]*]. For comparison of FHOD-1::GFP and mCherry::LEV-11E distributions in pharyngeal muscle, DWP275 and DWP 282 were each crossed with DWP3, resulting in strains DWP312 [*lev-11(ups175); qaIs8001[fhod-1::gfp]; upsEx178[myo-2p::mCH::lev-11E]*] and DWP313 [*lev-11(ups184); qaIs8001[fhod-1::gfp]; upsEx185[myo-2p::mCH::lev-11E]*], respectively. To examine the combined effects of loss of FHOD-1 and LEV-11E, DWP282 was crossed with XA8032, and progeny selected for loss of *him-5(e1461)* to generate DWP323 [*fhod-1(tm2363) lev-11(ups184); upsEx185[myo-2p::mCH::lev-11E]*].

### Muscle-enriched protein extraction

Worms were grown in the liquid culture as synchronized populations until the majority of animals were at least larval stages L3 and L4 for late-stage larval extracts, or as asynchronous populations for mixed-stage extracts. Worms were suspended in 30% sucrose and immediately centrifuged at 1500 x *g*, 5 min to separate lower-density live worms floating at the top from denser debris that pellets to the bottom. Collected worms were washed with ice-cold M9 buffer and 0.1 M NaCl solution before storage at −80°C overnight. Frozen worm pellets were thawed on ice in a cold room and resuspended in two volumes of ice-cold homogenization buffer (50 mM NaCl, 20mM Tris-HCl, pH 8.0, 1mM EDTA, 1mM DTT, 1mM PMSF), drop-frozen in liquid nitrogen, and pulverized into fine powder using a pre-chilled coffee grinder. The crude homogenate was subject to sonication on ice (30% amplitude, 50% duty cycle; twelve times 15 sec on/45 sec off, with 2 min off after every fourth run) before centrifugation for 10 min at 15,000 rpm, 4°C using a pre-chilled TLA 120.2 rotor. Pellets were washed twice with an equal volume of homogenization buffer to remove most actin monomers, free actin filaments, and non-body wall tissues. Actin and myosin filaments stably incorporated into myofibrils of body wall muscle were then extracted using an equal volume of extraction buffer (0.6 M KCl, 20 mM Tris-HCl, pH 8.0, 5 mM ATP, 5 mM MgCl_2_, 1 mM DTT, 1 mM PMSF, 0.2 mM EGTA). Extracted supernatant samples were clarified 10 min at 15,000 rpm, 4°C, and then flash frozen with liquid nitrogen and stored at −80°C.

### Immunoprecipitation mass spectrometry (IP-MS) analysis

200 µL of Affi-Prep Protein A beads (Bio-Rad, Hercules, CA) were washed four times with 1 mL of PBST (1xPBS, 0.1% Tween-20) and centrifuged at 6000 rpm (3500 x *g*), 30 sec before being resuspended in 1 mL of PBST buffer with an additional 100 µL (∼100 µg) of affinity-purified FHOD-1 antibody DPMSP2 (Mi-Mi et al., 2012) and incubated overnight at 4°C with agitation. The following day, beads were washed three times with 1 mL of PBST, three times with 1 mL of 0.2 M sodium borate (pH 9.0), then resuspended in 900 µL of 0.2 M sodium borate (pH 9.0) with 20 mM final concentration of fresh cross-linker, dimethyl-pimelimidate, and gently rotated 30 min at room temperature. The cross-linker was inactivated by washing beads with 1 mL of 0.2 M ethanolamine, 0.2 M NaCl (pH 8.5) twice before incubating with 1 mL of the same for 1 hr rotating, and storage in the same solution at 4°C overnight.

For mixed-stage and late-stage larvae extracts, two 100-µl aliquots of cross-linked FHOD-1 antibody-coupled beads were briefly washed 3 times with 1 mL of 0.5 M glycine (pH 2.5) to eliminate uncoupled antibody and reduce background precipitation, and then immediately washed three times with lysis buffer (100 mM KCl, 50 mM HEPES (pH 7.4), 1 mM EGTA, 1 mM MgCl_2_, 0.5 mM DTT, 10% Glycerol, 0.05% NP-40). Two immunoprecipitation reactions were performed in parallel by incubation of 75 µL of beads with 1 mL of muscle enriched extracts of DWP3 [*fhod-1(+)* with *fhod-1::gfp*] or XA8001 [*fhod-1(tm2363)*] worms for 5 hr at 4°C. Beads were then washed three times with 1 mL of lysis buffer with EDTA-free protease-inhibitor cocktail (Roche, Indianapolis, IN), and then three times with 1 mL of lysis buffer without NP-40 or protease-inhibitor cocktail. Immunoprecipitated proteins were eluted from beads with sample buffer without reducing agent, and afterwards supplemented with DTT to a final 100 mM and stored at −80°C until use.

Since SDS-PAGE tends to fail at resolving low abundance proteins or proteins with extreme molecular weights, the entire precipitation for each group as well as pre-immunoprecipitation muscle extracts of mixed stage worms were analyzed by LTQ-Orbitrap Tandem Mass Spectrometry (Han et al., 2008) at the Arizona Proteomics Consortium (University of Arizona, Tucson, AZ). Mass spectrometry data from the Arizona Proteomics Consortium were analyzed using Scaffold Viewer software (Version 5.3.2, Proteome Software Inc.). Minimum characteristics for protein identification confidence were 99.9% probability of correct protein, and 3 minimum peptides per protein with 95% probability of correct peptide identification.

### Antigen and antibody production and western blot analysis

To generate antibodies specific for short LEV-11 isoforms, guinea pigs (gp27-57, gp27-58) were immunized against keyhole limpet hemocyanin-conjugated peptide K-V-N-K-E-G-A-Q-Q-T-S-L-L-D-V-L-K-K-K, corresponding to amino acids 3-21 shared by all short LEV-11 isoforms but not long LEV-11 isoforms. To generate antibodies specific for long LEV-11 isoforms, chickens (ch10-42, ch10-43) were immunized against similarly conjugated peptide A-I-K-K-K-M-Q-A-M-K-I-E-K-D-N-A-L-D-R-C, corresponding to amino acids 3-22 of all long isoforms. Peptide conjugation and immunizations were performed by the Pocono Rabbit Farm and Laboratory, Inc. (Canadensis, PA).

For antibody affinity purification and characterization, LEV-11A, LEV-11C, and LEV-11E were produced as recombinant glutathione-S-transferase (GST) fusion proteins from *E. coli* bearing pGEX-6p-3-lev-11A, pLMM19, or pLMM25, respectively, and released from GST by cleavage with GST-3CPro Precision protease, as described previously (Amin et al., 2007). For affinity purification of antibodies for short LEV-11 isoforms, 2.5 mg LEV-11C was resolved by SDS-PAGE and stained in-gel 3 min with Coommassie brilliant blue to allow excision of LEV-11C-containing portions of polyacrylamide gel. LEV-11C was electroeluted from gel portions into coupling buffer (0.1 M NaHCO_3_, pH 8.3, 0.5 M NaCl). CNBr-Sepharose beads (1 g) were hydrated in water for 10 min before activation in 40 ml of 10 mM HCl, followed by further continuous wash with 210 ml of 10 mM HCl over a sintered glass filter, and a final wash with 40 ml of ice-cold coupling buffer. The activated and washed CNBr-Sepharose beads were resuspended in 8 ml of coupling buffer containing electroeluted LEV-11C, and mixed 2 hr at room temperature to allow coupling. LEV-11C-coated beads were then incubated over night at 4°C in 0.2 M glycine, pH 8.0 to quench further CNBr-reactivity. Beads were then washed with 20 ml of ice-cold acetate buffer (0.1 M Na-acetate, pH 4.0, 0.5 M NaCl) followed by 20 ml of ice-cold coupling buffer. This sequence of washes was repeated four additional times. LEV-11C-coated beads were then suspended in 2 ml of gp27-57 crude antiserum with 5 mM NaN_3_ overnight at 4°C. Antibody-coated beads were washed with 40 mL of ice-cold coupling buffer, and then eluted with 6 mL of 0.1 M glycine-HCl, pH 2.5 as 500-µL fractions that were each immediately neutralized with 120 µL of Tris-HCl, pH 8.5. Fractions 4-8 were pooled as ∼ 0.2 mg/ml affinity purified anti-LEV-11(Short) DPMSP13. These were supplemented to 5 mM NaN_3_ and 2 mg/ml BSA for storage at −80°C. LEV-11C-coupled beads were stored at 4°C in coupling buffer with 5 mM NaN_3_.

Western blot analysis (Fig.2) was conducted as previously published (Yingling & Pruyne, 2021). Sample sizes were normalized based on whole protein Coomassie brilliant blue stain intensity after SDS-PAGE. Affinity purified anti-LEV-11(Short) DPMSP13 and crude anti-LEV-11(Long) serum ch10-42 were diluted 1:200 in 1% milk/TBS/0.1% Tween 20 and incubated with primary antibody for 2 hr at room temperature. Images were acquired using a Bio-Rad ChemiDoc MP imager, and processed with Image Lab and Photoshop CS4.

### Fluorescence Microscopy

F-actin staining of worms (Figures 4B, 6, 7A) was done as previously published (Mi-Mi et al., 2012). Immunostain (Fig.4A, 7B, 7B, 8C) was done based on prior protocols (Finney & Ruvkun, 1990), but without the addition of spermidine-HCl and with a methanol concentration of 75% during fixation. Monoclonal antibody MH35 (anti-ATN-1) was developed by R.H. Waterston (Francis & Waterston, 1985) and generously provided as a gift by P. Hoppe (Western Michigan University, Kalamazoo, MI), and rabbit anti-GFP was a gift from Anthony Bretscher (Cornell University, Ithaca, NY). Primary antibody dilutions were MH35 1:10^4^, DPMSP13 affinity-pure anti-LEV-11(Short) 1:10, ch10-42 crude anti-LEV-11(Long) serum 1:100, anti-GFP 1:200, anti-mCherry 1:1000 (#632543 Takara Bio, San Jose, CA). Secondary antibodies donkey anti-Guinea pig Texas Red and goat anti-mouse FITC (Rockland Immunochemicals, Limerick, PA) were diluted 1:500.

Confocal images were acquired with an SP8 Laser Scanning Confocal Microscope (Leica, Wetzlar, Germany) driven by LAS X software (Version 3.5.2, build 4758; Leica) with a HCX Plan Apochromat 63X/NA 1.4 oil immersion lambda objective. Z-stack images were collected at 0.3 µm intervals for body wall muscle images and 0.2 µm for pharyngeal images. Images were deconvolved with Huygens Essential Software (Huygens compute engine 18.10.0, Scientific Volume Imaging B.V.), Classic Maximum Likelihood Estimation deconvolution algorithm, with 40 iterations and a signal to noise ratio of 20. Images were linearly processed to enhance contrast and false colored in Adobe Photoshop CS4 or CC2018 (Adobe, San Jose, CA). Deconvolved confocal z-stacks were used to generate X, Y, Z plane pharyngeal images using Imaris x64 software (version 10.1.0; Bitplane AG, Belfast, United Kingdom).

For analysis of mosaic pharynges (Fig.3E), age synchronized animals were fixed and mounted on slides. Pharynges were considered mosaic if at least one but not all of the major pharyngeal muscles (pm3-7) visibly express mCherry for DWP 272 [*lev-11(ETU-); Ex[mCH::lev-11E]*] n = 86 L1 larvae, n= 70 day-1 adults and for DWP282 [*lev-11(CETU-); Ex[mCH::lev-11E]*] n = 105 L1 larvae, n = 75 day-1 adults.

Analyses of BWM cell width and striations (Fig.8A,B) were done as previously published (Kimmich et al., 2024). Age synchronized day-1 adult N2, XA8001 [*fhod-1(*Δ)], DWP 262 [*lev-11(C-)*], DWP 263[*lev-11(C-)*], and DWP264[*lev-11(C-)*] were fixed and stained with Alexa568-Phalloidin. 15-20 worms were analyzed per strain with two to four muscle cells measured per worm (one muscle cell per muscle). For consistency, only muscle cells within two to four muscle cells from the vulva were analyzed.

### Pharyngeal pumping analysis

L1 larvae were placed on an NGM plate containing OP50-1. Using a Nikon SMZ18 fluorescence stereoscope (Nikon, Tokyo, Japan) equipped with a Sola light engine (Lumencore, Beaverton, OR), presence or absence of pharyngeal pumping in individual worms was determined by visual observation of the grinder bulb of the pharynx, followed by examination of the pharyngeal muscle under fluorescence illumination for mCherry expression (Fig.3F). 50-60 larvae were analyzed per strain in each of three independent trials. Movies of pharyngeal pumping (Movie 1, Movie 2) were captured using a Canon EOS Rebel 600D T3i camera with a Martin Microscope Company MM-SLR Adapter on a Nikon SMZ18 stereoscope. Movie files were edited and processed using iMovie (version 10.4.2, Apple, Cupertino, California).

### Viability analysis

To analyze viability (Fig.3A, C, D), young adult worms were set onto separate fresh NGM/OP50-1 plates and allowed to lay eggs for 3 hr. Eggs were counted on each plate. The next day (day 2), unhatched eggs (indicative of failed embryogenesis) and L1 stage larvae were counted. On days 2 - 4, viable worms were counted. For wild type N2 worms and strains for which we could isolate homozygous mutants, this analysis was performed with ten initial worms. For strains that we were unable to isolate homozygous animals (DWP267 and DWP280), analysis was performed with 40 adult progeny of heterozygous animals that, after the initial egg laying, were subject to single worm PCR to genotype and identify wild-type and heterozygous animals.

### Brood Size Analysis

To measure brood sizes (Fig.3B), ten L4 animals were set onto separate NGM/OP50-1 plates and allowed to lay eggs. Every 12 hr, worms were moved to fresh plates, and eggs were counted. This was repeated until 5 days had elapsed.

### Statistical Analysis

Analysis of variance was performed on data containing three or more groups. Unpaired T-tests were performed on comparisons of two groups. Differences for which *p* < 0.05 were considered statistically significant. Graphs were made and post hoc analyses conducted using Prism 10 (version 10.1.1; GraphPad Software, Boston, MA) with data presented as average ± standard deviation.

## ACKNOWLEDGMENTS

Thaks to WormBase and WormAtlas. Some worm strains were generated in collaboration with InVivo Biosystems. Other worm strains were obtained from the CGC, which is supported by NIH Office of Research Infrastructure Programs (P40 OD010440). This work was supported by National Institute of Arthritis and Musculoskeletal and Skin Diseases of the NIH under Award Number R01AR064760 to D.P. and the Fredrick Hendricks Endowment Fund under Award Number 102234 to D.P.

---

**Movie 1. mCH::LEV-11E expression in pharyngeal muscle supports pharyngeal pumping in mutants containing no endogenous short LEV-11 isoforms.** A *lev-11 (CETU-)* L1 larva confirmed to express mCH::LEV-11E in its pharynx exhibits visible pharyngeal pumping (arrow) as it crawls among bacteria (food).

**Movie 2. Absence of short LEV-11 isoforms impedes pharyngeal pumping.** A *lev-11 (CETU-)* L1 larvae confirmed to not express mCH::LEV-11E in its pharynx exhibits no pumping of the pharynx (arrow) as it moves on a plate containing bacteria (food).

## SUPPORTING INFORMATION

**Supplementary Table S1. Total spectrum counts for proteins in pre-IP extracts of mixed stage animals.**

**Supplementary Table S2. Total spectrum counts for proteins in FHOD-1 IP of mixed stage extracts.**

**Supplementary Table S3. Total spectrum counts for proteins in FHOD-1 IP of late larval stage extracts.**

**Supplementary Table S4. Worm strains used in this study.**

